# Ensemble cryo-electron microscopy reveals conformational states of the nsp13 helicase in the SARS-CoV-2 helicase replication-transcription complex

**DOI:** 10.1101/2021.11.10.468168

**Authors:** James Chen, Qi Wang, Brandon Malone, Eliza Llewellyn, Yakov Pechersky, Kashyap Maruthi, Ed T. Eng, Jason K. Perry, Elizabeth A. Campbell, David E. Shaw, Seth A. Darst

**Author notes:** These authors contributed equally to this work (James Chen, Qi Wang, Brandon Malone).

## Abstract

The SARS-CoV-2 nonstructural proteins coordinate genome replication and gene expression. Structural analyses revealed the basis for coupling of the essential nsp13 helicase with the RNA dependent RNA polymerase (RdRp) where the holo-RdRp and RNA substrate (the replication-transcription complex, or RTC) associated with two copies of nsp13 (nsp13_2_-RTC). One copy of nsp13 interacts with the template RNA in an opposing polarity to the RdRp and is envisaged to drive the RdRp backwards on the RNA template (backtracking), prompting questions as to how the RdRp can efficiently synthesize RNA in the presence of nsp13. Here, we use cryo-electron microscopy and molecular dynamics simulations to analyze the nsp13_2_-RTC, revealing four distinct conformational states of the helicases. The results suggest a mechanism for the nsp13_2_-RTC to turn backtracking on and off, using an allosteric mechanism to switch between RNA synthesis or backtracking in response to stimuli at the RdRp active site.

---

COVID-19, caused by the coronavirus SARS-CoV-2 ^1,2^, continues to devastate the world. The viral RNA-dependent RNA polymerase (RdRp, encoded by non-structural protein 12, or nsp12) functions as a holo-RdRp (comprising nsp7/nsp8_2_/nsp12) in a replication-transcription complex (holo-RdRp + RNA, or RTC) to direct RNA synthesis from the viral RNA genome ^3–5^. The RdRp is also a target for the clinically approved antiviral remdesivir ^6–8^. In addition to the holo-RdRp, the virus encodes several nucleic acid processing enzymes, including a helicase (nsp13), an exonuclease (nsp14), an endonuclease (nsp15), and methyltransferases (nsp14 and nsp16) ^9^. Little is known about how these enzymes coordinate to replicate and transcribe the viral genome.

Nsp13, essential for viral replication ^10–13^, is a superfamily 1B (SF1B) helicase that can unwind DNA or RNA substrates with a 5′->3′ directionality ^14–16^. Along with the two canonical RecA ATPase domains of SF1 helicases ^14,17^, nsp13 contains three additional domains; an N-terminal zinc-binding domain (ZBD, unique to nidoviral helicases), a stalk, and a 1B domain ^13,18,19^. Prior studies established that the nsp13 helicase forms a stable complex with the RTC, and single-particle cryo-electron microscopy (cryo-EM) structures of an nsp13_2_-RTC (the RTC with two nsp13 protomers bound) have been determined ^20–22^.

In the nsp13_2_-RTC structure, two protomers of nsp13 (nsp13.1 and nsp13.2; Fig. 1) sit on top of the RTC with each nsp13-ZBD interacting with one of the two N-terminal helical extensions of nsp8 ^20–22^. This overall architecture places the nsp13.1 active site directly in the path of the downstream template-RNA (t-RNA). The cryo-EM maps showed that the 5′-single-stranded overhang of the t-RNA (Fig. S1) passed through the nucleic acid binding channel of nsp13.1 ^23^, but the low resolution of the map due to structural heterogeneity precluded detailed modeling ^20^.

**Fig. 1.**
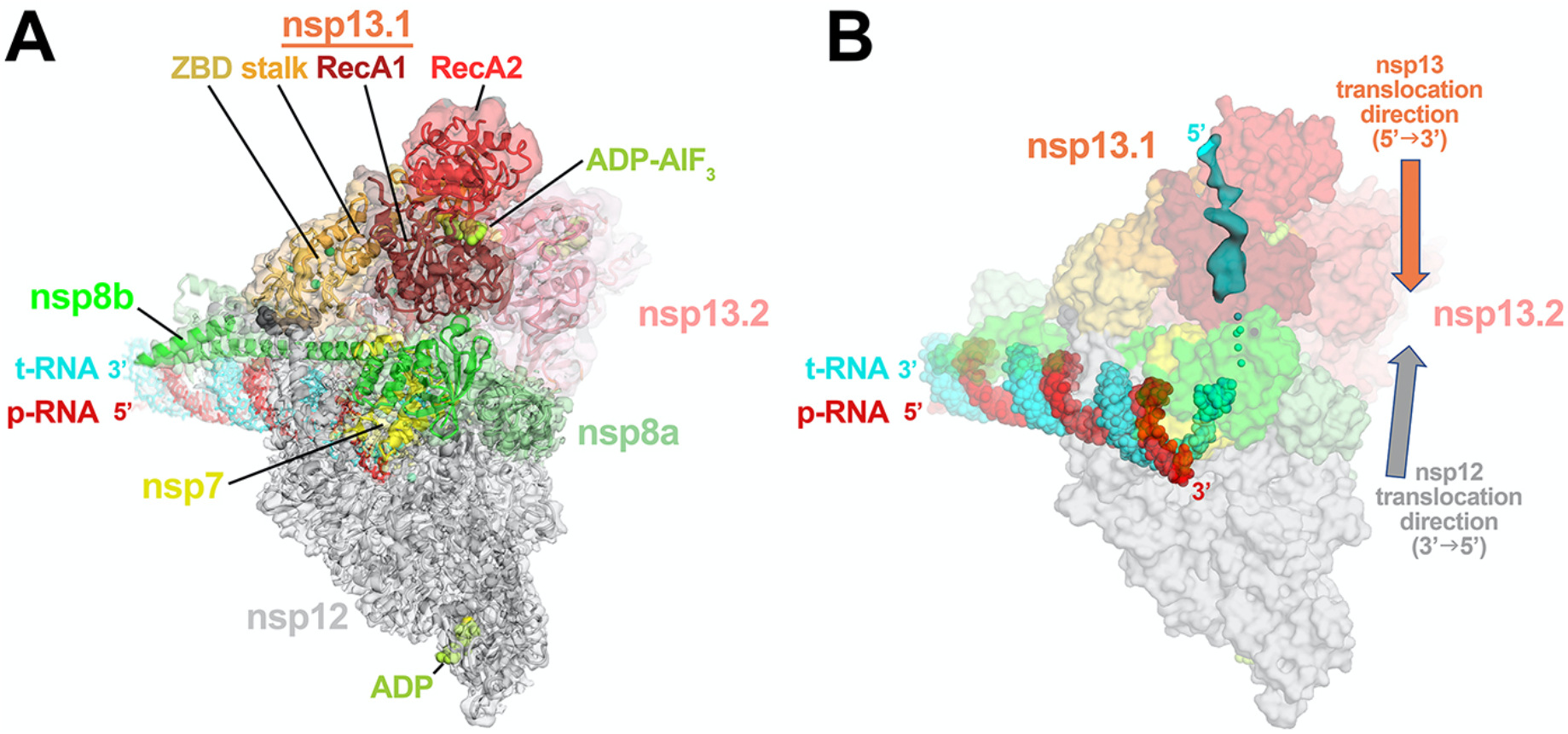
Consensus cryo-EM structure of an nsp13_2_-RTC. **A.** Overall architecture of the consensus nsp13_2_-RTC. Shown is the transparent cryo-EM density (map3, local-resolution filtered) with the nsp13_2_-RTC model superimposed. **B.** The consensus nsp13_2_-RTC structure is shown; RNA is shown as atomic spheres, proteins are shown as transparent molecular surfaces. A low-pass filtered (6 Å) cryo-EM difference density reveals the path of the downstream t-RNA 5′-segment through the RNA binding groove of nsp13.1 (cyan surface). See also Figs. S1-S4.

The structural analysis of the nsp13_2_-RTC provided new perspectives into the role of the nsp13 helicase in the complex viral replication-transcription program, suggesting that nsp13 may facilitate processive elongation by the RdRp on the highly structured RNA genome ^24,25^, but may also generate backtracked RTCs for proofreading, template-switching during sub-genomic RNA transcription, or both ^20,26^. How nsp13 directs these diverse processes that regulate RdRp function remains less understood. For instance, the structures indicate that nsp13 translocates on the t-RNA strand in the 5′->3′ direction ^16^, while the RdRp would translocate on the same strand in the opposite direction (Fig. 1B). How can the RdRp rapidly replicate the ~30 kb viral genome ^27^ if it is opposed by the helicase? Also, what is the role of the second nsp13 protomer (nsp13.2), which appears capable of ATPase and translocation/helicase activity but does not appear to be engaged with nucleic acid in the structures ^20,26^?

Here we describe an extensive structural analysis of a cryo-EM dataset of the nsp13-RTC, combined with molecular dynamics (MD) simulation analysis of the resulting structures. The results yield a cryo-EM map of the nsp13_2_-RTC at a nominal resolution of 2.8 Å (2.1-2.5 Å in the active site core of the RdRp; Fig. 1). Structural heterogeneity apparent in the nsp13 portions of the map was resolved by classification approaches, revealing four distinct conformational states of the nsp13 subunits. Analysis of these conformational states suggest solutions to the apparent contradictions regarding the role of nsp13 and provides further insight into models for nsp13 function during viral replication/transcription.

## RESULTS

### An augmented cryo-EM dataset allows extensive structural analysis of the nsp13-RTC

Previously we described a single-particle cryo-EM analysis of a stable SARS-CoV-2 nsp13-RTC from a curated set of 88,058 particle images ^20^. These particles were classified into three distinct assemblies, nsp13_1_-RTC (4.0 Å nominal resolution), nsp13_2_-RTC (3.5 Å), and a dimer of nsp13_2_-RTC [(nsp13_2_-RTC)_2_; 7.9 Å]. Here we analyzed a much larger dataset (nearly five times as many particles; Fig. S1, Table S1) collected from the same sample preparation. From a consensus refinement (Figs. S1 and S2, map1; Note: Fig. S1 shows the details of the cryo-EM processing pipeline; Fig. S2 is a streamlined cryo-EM processing pipeline that highlights the essential steps), the particles were classified ^28^ into the same three assemblies observed previously [nsp13_1_-RTC (map2), nsp13_2_-RTC (map3), (nsp13_2_-RTC)_2_ (map4)] ^20^ with a very similar distribution of particles between the three assemblies (Figs. S1 and S2; Tables S1 and S2), confirming the robustness of the classification procedure. We focus primarily on the nsp13_2_-RTC because the bulk of the particles (72%) belong to this class and generated the highest resolution map (Figs. S1, S2 and S3; map3; 2.9 Å nominal resolution).

To obtain the best possible consensus cryo-EM map of the entire complex, we generated a series of cryo-EM maps by focused refinement around sub-domains of the nsp13_1_-RTC (map2) and nsp13_2_-RTC (map3) maps and combined these, generating a composite map with a nominal resolution of 2.8 Å (Fig. 1; Figs. S1-S3, map9). Local resolution analysis ^29^ suggested that the active site and NiRAN ligand-binding pocket of the RdRp were resolved to between 2.1-2.6 Å resolution (Fig. S3). This was supported by the excellent quality of the cryo-EM map, where the ADP-Mg^2+^ bound in the NiRAN domain enzymatic site could be visualized (Fig. S4), and RNA base pairs near the RdRp active site could be identified directly from the cryo-EM density (Fig. S4). Although not as well resolved, the ADP-AlF_3_-Mg^2+^ and surrounding residues in the nsp13 active sites could also be modeled (Fig. S4).

Despite the excellent map quality for most of the RTC (Figs. 1 and S4), features of the composite consensus map (map9) suggested substantial heterogeneity in the nsp13 subunits, particularly in the RecA2 and 1B domains (Fig. S3). Therefore, we generated a mask surrounding the nsp13.1 and nsp13.2 RecA1, RecA2, and 1B domains (of map3; Figs. 2 and S2) and used masked classification with signal subtraction ^30^ to identify four distinct conformational states (Figs. 2, S1, S2, and S5; Table S3) with significant differences in the dispositions of the nsp13 subunits, particularly nsp13.1.

**Fig. 2.**
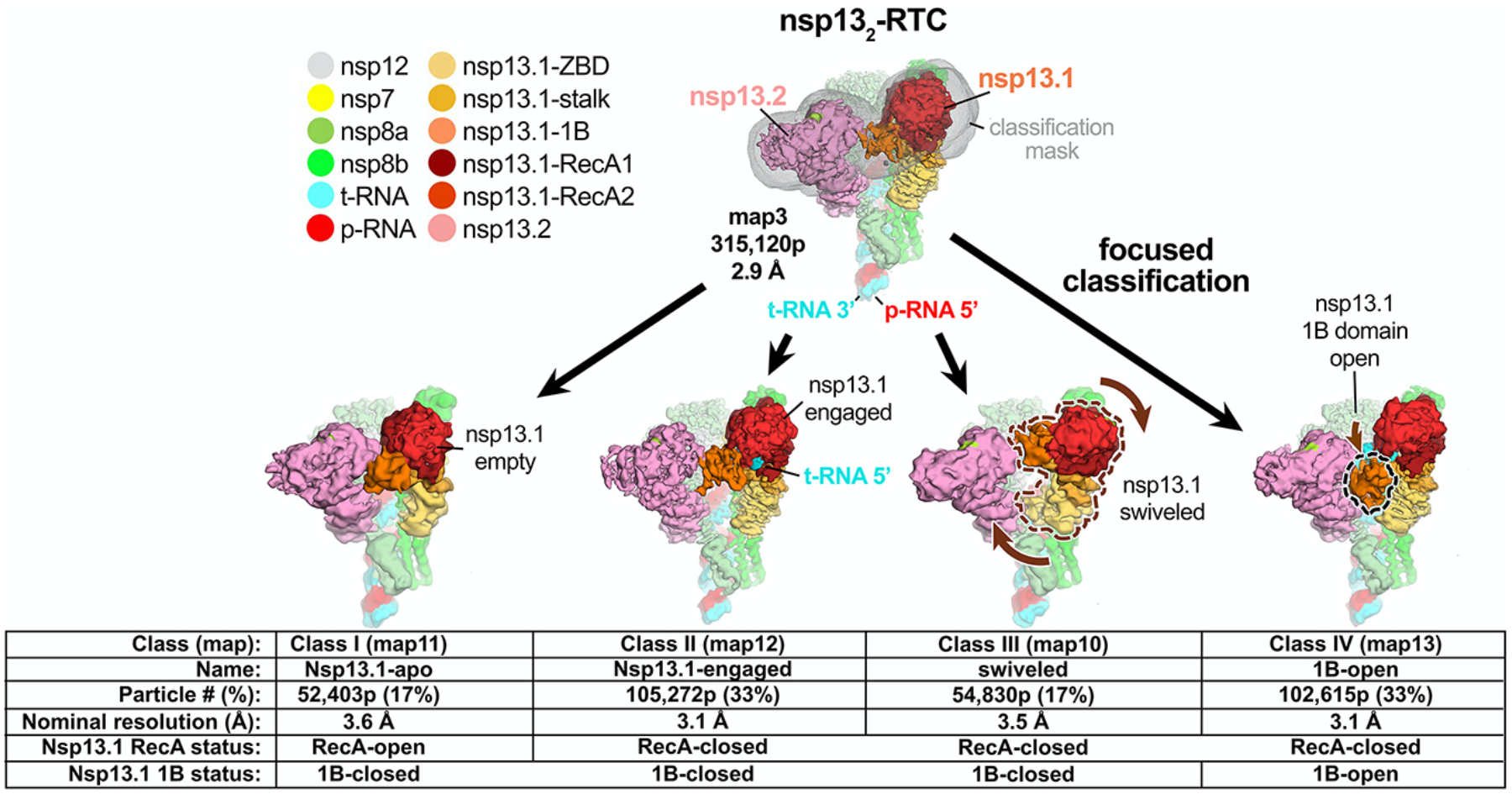
Four conformational states of the nsp13_2_-RTC. (*top*) Cryo-EM density (map3, local-resolution filtered) colored according to the code on the left. A mask was constructed surrounding the nsp13.1 and nsp13.2 1B, RecA1, and RecA2 domains (grey mesh). The 315,120 particles were divided into four distinct structures (class I, II, III, and IV) by focused classification inside the mask, followed by further refinement (Figs. S1, S5). Class II contained the most particles, and the nsp13.1 RecA domains were completely closed (Fig. S5), entrapping the 5′-t-RNA segment in a groove between the two RecA domains and the 1B domain (Fig. 3). Therefore, class II (nsp13.1-engaged) was used as a reference for comparison of the other structures. Each class was characterized by one dominant conformational change: class I) nsp13.1-apo, the RecA domains were completely open (Fig. S5) and devoid of RNA (Fig. 4), class III) swiveled, the nsp13.1 protomer as a whole was rotated 38° as shown (Fig. 6), class IV) 1B-open, the nsp13.1 1B domain was rotated open by 85° (Fig. 5). Also see Fig. S5 and Videos S1 and S2.

The class II structure (Figs. 2 and S5) contains the most particles and the nsp13 subunits are best resolved in this map (map12; Figs. 2, S2 and S5). Compared to the other structures, the nsp13.1 RecA domains of class II (map12) are closed onto each other more than the other structures (Fig. S5) and are thereby engaged most tightly with the RNA (see below). We call this the ‘nsp13.1-engaged’ structure and use it as a reference to give a general overview of the conformational changes in the other classes.

While each of the classes shows significant changes in both the disposition of each nsp13 subunit as a whole as well as intramolecular domain motions within each nsp13 subunit, each structural class can be characterized by one dominant conformational change in nsp13.1 (compared to the nsp13.1-engaged structure used as a reference):

i. In class I, the nsp13.1 RecA2 domain is rotated open by 21° with respect to RecA1. Concomitantly, the RNA binding site is empty while occupancy of the nsp13.1 nucleotide-binding site is ambiguous. We therefore call this the ‘nsp13.1-apo’ structure (Fig. 2).
ii. In class III, the nsp13.1 subunit swivels as a whole by 38° away from nsp13.2. We call this the ‘nsp13.1-swiveled’ structure (Fig. 2).
iii. In class IV, the nsp13.1 domain 1B is rotated 85° away from the nsp13.1 RNA binding channel, creating the ‘1B-open’ structure (Fig. 2).

### The nsp13.1-engaged conformation grasps the downstream RNA t-strand

In the nsp13.1-engaged structure, the distance between the center-of-gravity of the two nsp13.1 RecA domains, 27.3 Å, is the shortest of the eight nsp13 conformations (Fig. S5). The RecA domains are thus ‘closed’ and grasp the downstream t-RNA single-stranded 5′-segment emerging from the RdRp active site, giving rise to well-resolved cryo-EM density for the RNA passing through the helicase (Fig. 3A). The RNA is corralled in a tunnel between the two RecA domains and the 1B domain, which is also in a closed conformation (Figs. 2 and 3A). The pattern of purine and pyrimidines in the cryo-EM density is clearly discernable, allowing the unique sequence register of the RNA engaged with the nsp13.1 helicase to be determined (Fig. 3A).

**Fig. 3.**
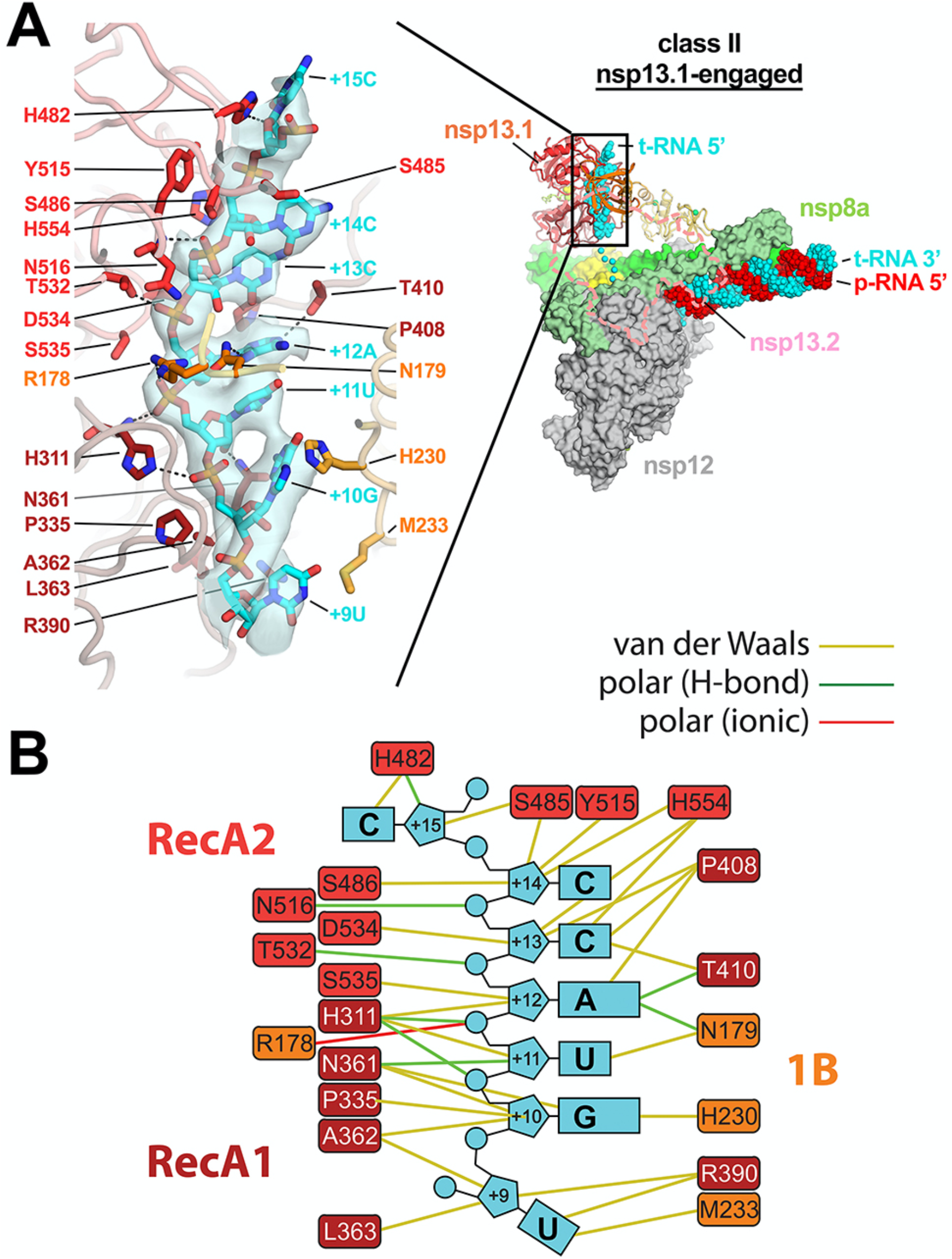
In class II (nsp13.1-engaged), the nsp13.1 RecA domains and 1B domain clamp onto the 5′-single-stranded t-RNA. **A.** (*right*) Overall view of the nsp13.1-engaged structure. Proteins are shown as molecular surfaces except nsp13.1 is shown as a backbone ribbon, and nsp13.2 is removed and shown only as a dashed outlline. The RNA is shown as atomic spheres. The boxed region is magnified on the left. (*left*) Nsp13.1 is shown as a backbone worm but with side chains that interact with the t-RNA shown. Cryo-EM density for the downstream 5′-t-RNA segment is shown (transparent blue surface) with the t-RNA model superimposed. The pattern of purines/pyrimidines in the RNA density was clear and unique, allowing the identification of the sequence register for the nsp13.1-bound RNA. **B.** Schematic illustrating nsp13.1-RNA interactions. Also see Video S2.

The ordered RNA segment is 7 nucleotides in length (+9 to +15; Fig. 3), with the five central nucleotides (+10 to +14) completely enclosed within the helicase. The RNA phosphate backbone generally faces the nsp13.1 RecA domains, and the mostly stacked bases face the 1B domain (Fig. 3). As might be expected, the helicase establishes extensive interactions with the RNA phosphate backbone, including several polar interactions. Interactions with the RNA bases are mostly van der Waals interactions and not expected to be base-specific (Fig. 3).

### The nsp13.1-apo state

Comparison of the nsp13.1-apo and nsp13.1-engaged structures revealed a striking change in the conformation of the RecA-like ATPase domains of nsp13.1. Superimposition of the α-carbons of nsp13.1 RecA1 (residues 235-439) or RecA2 (residues 440-596) alone yielded root-mean-square-deviations (rmsds) of 0.387 and 0.673 Å, respectively, indicating the structures of the individual domains are very similar between the two states. However, superimposition via the α-carbons of only RecA1 gave an rmsd of 7.05 Å for the RecA2 α-carbons, indicating a substantial change in the relative disposition of the two domains. The movement of RecA2 with respect to RecA1 corresponds to an ~21° rotation about the axis shown in Fig. 4A (also see Video S1), corresponding to an opening of the RecA domains; the nsp13.1 RecA domains of the nsp13.1-apo state are the furthest open of any of the eight nsp13 protomer structures (Fig. S5F).

**Fig. 4.**
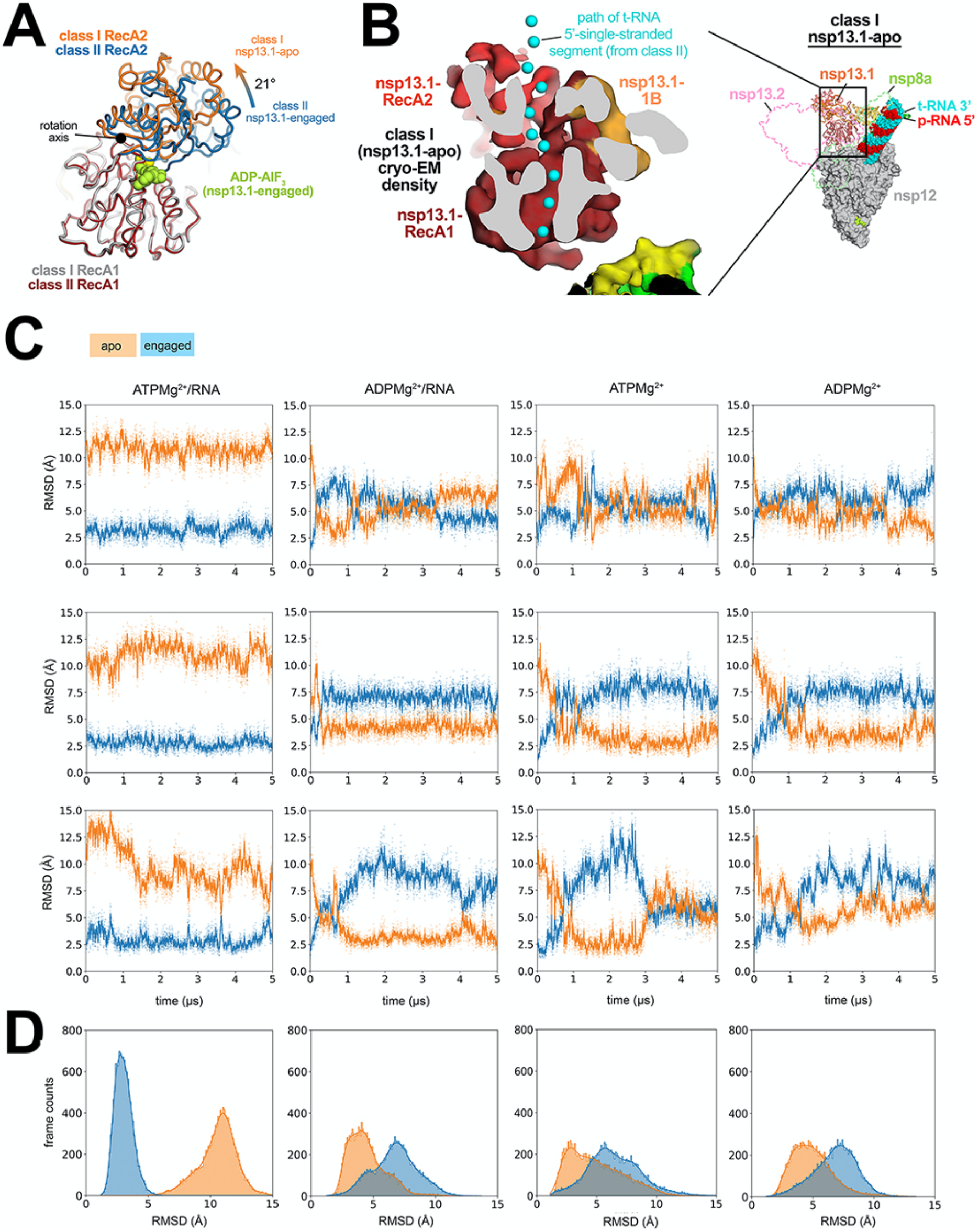
In nsp13.1-apo, the RecA domains are open and devoid of RNA density. **A.** The RecA1 domains of class I (nsp13.1-apo) and class II (nsp13.1-engaged) were superimposed (rmsd of 0.387 Å over 205 α-carbons), revealing that the RecA2 domain of nsp13.1-apo was rotated open by 21° about the rotation axis shown. The ADP-AlF_3_-Mg^2+^ from the nsp13.1-engaged structure is shown as limon atomic spheres. **B.** (*right*) Overall view of the nsp13.1-apo structure. Proteins are shown as molecular surfaces except nsp13.1, which is shown as a backbone ribbon, and nsp13.2, which is removed and shown only as a dashed outline. The RNA is shown as atomic spheres. The boxed region is magnified on the left. (*left*) Cryo-EM density of nsp13.1-apo reveals that the RNA path is empty (the RNA path from the nsp13.1-engaged structure is denoted by cyan spheres). **C.** Three independent simulations of nsp13.1 bound to ATPMg^2+^/RNA, ADPMg^2+^/RNA, ATP Mg^2+^, and ADPMg^2+^. Values of rmsd plotted represent the heavy-atom rmsd of the RecA2 lobe (after alignment on the RecA1 lobe) with respect to nsp13.1-engaged (blue) and nsp13.1-apo (orange) cryo-EM structures. **D.** The rmsd histograms represent aggregate values across all three replicates shown in (**C**). Also see Video S1.

The consensus nsp13_2_-RTC cryo-EM map (map3; Figs. S1 and S2) contains low-resolution density indicating that the downstream single-stranded 5′-segment of the t-RNA occupies the nsp13.1 RNA binding channel (Fig. 1B). Moreover, the t-RNA 5′-segment occupying the nsp13.1 RNA binding channel of the nsp13.1-engaged state is well resolved (Fig. 3). By contrast, the nsp13.1-apo cryo-EM density shows that the nsp13.1 RNA-binding path is empty (Fig. 4B). The nsp13.1-apo cryo-EM density also does not support occupancy of ADP-AlF_3_-Mg^2+^ in the nucleotide-binding site of nsp13.1, although the low resolution of the map in this region makes this conclusion tentative.

### Spontaneous and reversible transition of the nsp13.1 RecA domains between the nsp13.1-engaged and nsp13.1-apo conformations

To characterize the RecA1-RecA2 interdomain movement and how a bound substrate may influence that movement, we performed MD simulations of free nsp13.1 (i.e., without nsp13.2 or the RTC) under four different substrate-bound conditions (ATPMg^2+^/RNA, ADPMg^2+^/RNA, ATPMg^2+^ only, and ADPMg^2+^ only). For each condition, we ran three independent 5-μs simulations, all initiated from the nsp13.1-engaged conformation (Figs. 2 and 3).

In simulations of ATPMg^2+^/RNA-bound nsp13.1, the RecA2 domain maintained its general orientation with respect to RecA1 throughout the simulations (Fig. 4C). The average rmsd of RecA2 between the initial nsp13.1-engaged cryo-EM structure and the structures from the MD trajectories, aligned on the RecA1, was low (∼2.9 Å; some adjustment of RecA2 from the initial nsp13.1-engaged cryo-EM structure position in these simulations was expected, as the cryo-EM structure was determined using ADP-AlF_3_/RNA in place of ATPMg^2+^/RNA). Conformations resembling the nsp13.1-apo structure (rmsd <3.5 Å) were not observed (Fig. 4C, D).

In simulations of ADPMg^2+^/RNA-bound nsp13.1, RecA2 rotated away from its initial position in the nsp13.1-engaged conformation, and nsp13.1-apo–like conformations were repeatedly visited throughout the simulations (Figs. 4C, D). The ADPMg^2+^/RNA-bound nsp13.1-apo-like conformations were metastable, and interconverted with the nsp13.1-engaged conformations. Spontaneous and reversible conversion between the nsp13.1-engaged and nsp13.1-apo conformations was also observed in the simulations of ATPMg^2+^-bound and ADPMg^2+^-bound nsp13.1 (Figs. 4C, D). These results suggest that the presence of both the ATPMg^2+^ and RNA may stabilize the nsp13.1-engaged conformation and that the absence of either substrate may destabilize the nsp13.1-engaged conformation and facilitate the transition to the nsp13.1-apo conformation, consistent with the observations from the cryo-EM analysis.

### The ‘1B-open’ conformation of nsp13.1 may explain how the RdRp can synthesize RNA in the presence of nsp13

In the nsp13.1-engaged state, the downstream single-stranded t-RNA is guided through a deep groove between the RecA1 and RecA2 domains that is completely closed off by the 1B domain (Fig. 5A). Remarkably, in the 1B-open structure, the nsp13.1 1B domain rotates 85° about the stalk away from the nsp13.1 RNA binding channel, creating an open groove rather than a closed tunnel (Fig. 5B). The cryo-EM density allows modeling of the downstream single-stranded t-RNA emerging from the RdRp active site up to the edge of the open groove proximal to the RdRp, but the RNA density disappears there, indicating that the RNA is not engaged within the active site of the helicase (Fig. 5B).

**Fig. 5.**
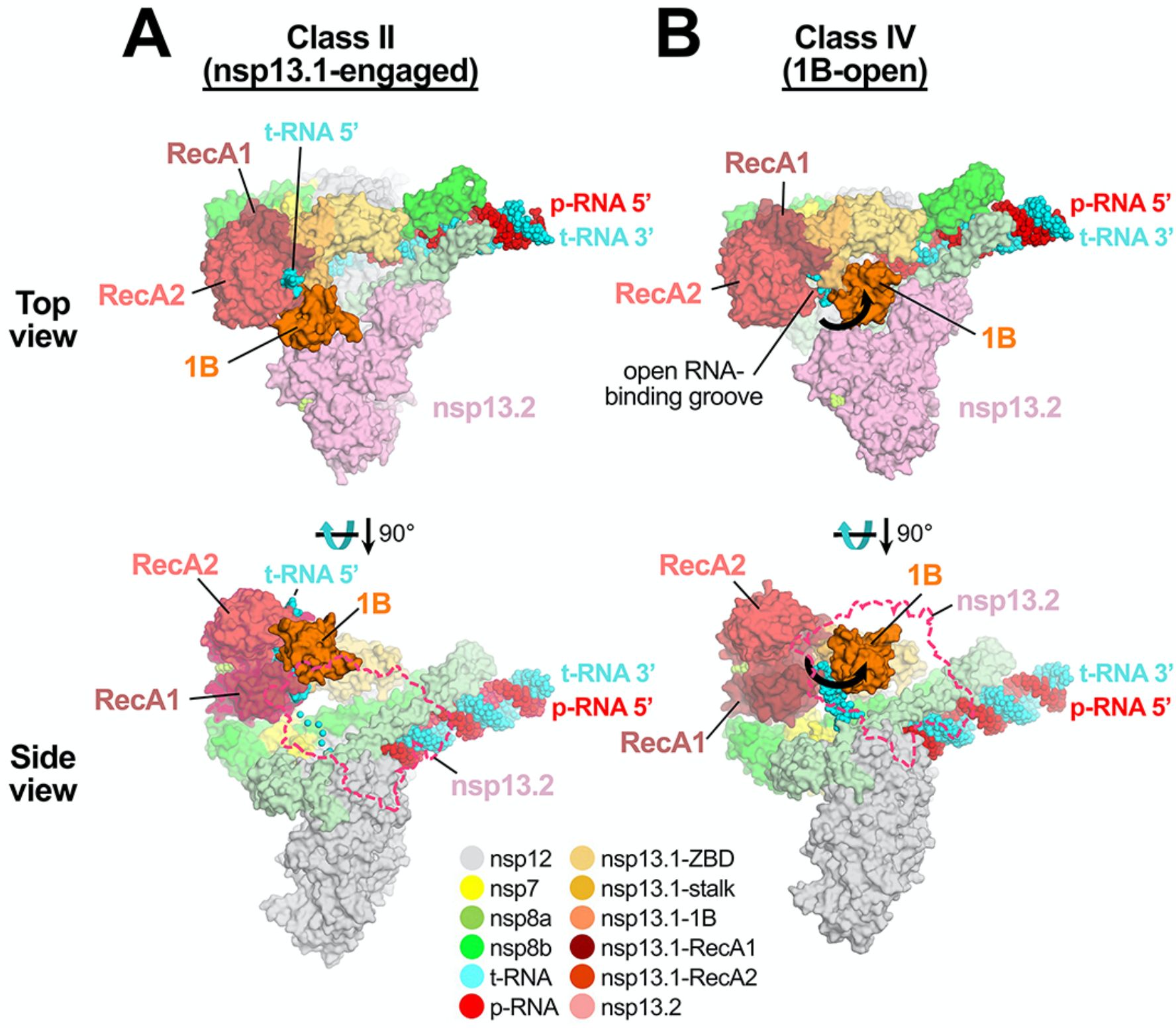
1B-open structure. Comparison of nsp13.1-engaged (A) and 1B-open (B) structures. Two views are shown, a top view (*top*) and a side view (*bottom*). In the top view, the proteins are shown as molecular surfaces and color-coded according to the key at the bottom. In the side view, nsp13.2 is shown only as a dashed outline. The RNA is shown as atomic spheres. In the 1B-open structure (B), the nsp13.1 1B domain is rotated open by 85° (represented by thick black arrows). The 5′-t-RNA emerging from the RdRp active site approaches the nsp13.1 RNA binding groove but does not enter it. Also see Video S2.

In the 1B-open conformation, the nsp13.1 1B domain appears to be trapped open by the presence of nsp13.2 (Fig. 5B), with the transition from the 1B-open to the 1B-closed conformation blocked by nsp13.2. Consistent with this, we analyzed the nsp13 conformational states in the nsp13_1_-RTC (nsp13.2 absent) by masked classification with signal subtraction around the RecA1, RecA2 and 1B domains of the single nsp13 (Fig. S1) but the 1B-open nsp13 conformation was not observed. We propose that the 1B-open conformation of the nsp13.1 1B domain is trapped by the presence of nsp13.2.

We note that in the (nsp13_2_-RTC)_2_ dimer (Figs. S1-S3), the nsp13 protomers corresponding to nsp13.1 are also in the ‘1B-open’ state, as was observed by Yan *et al.*^31^. Since the dimer only comprises 8% of our particle dataset (Table S1) while the nsp13_2_-RTC complex comprises 72% of the particles, we have focused our attention on the latter complex. We observe that the (nsp13_2_-RTC)_2_ dimer forms in the absence of additional factors such as nsp10-14 ^20^, in contrast to what’s reported in Yan *et al.* ^31^.

Yan *et al.* ^31^ observed the 1B-open state of nsp13.1 (labeled nsp13-2 in their nomenclature) in their (dimer) dCap(0)-RTC structure, curiously assigned as a backtracking-competent state. This is at odds with: i) observations that nsp13.1 in the 1B-open conformation does not engage RNA in its RNA-binding groove [Fig. 5B; also observed by Yan *et el.* ^31^] and so would be incompetent for RNA translocation, and ii) the finding that nsp13 stimulated SARS-CoV-2 RTC backtracking ^26^.

### Spontaneous transition of the nsp13.1 1B domain from the 1B-open to 1B-closed conformations

The conformations of the nsp13.1 1B domain in the nsp13.1-engaged and nsp13.1-apo structures are related by a ~10° rotation around the nsp13-stalk, but the 1B domains are closed on the nsp13-RecA domains in both structures. We refer to these collectively as ‘1B-closed’ states (Fig. 2). These conformations have also been observed in crystal structures of isolated nsp13 as well as some other SF1-like helicases ^32^. The conformation of the 1B domain in the 1B-open cryo-EM structure, in which the domain is rotated ~85° compared to the 1B-closed conformations, was only seen in nsp13.1 when it was paired with nsp13.2 in the RTC, suggesting that this conformation may not be stable in isolated nsp13. To test this hypothesis, we performed five independent 25-μs simulations on isolated (free) nsp13 (with ADPMg^2+^), initiated from the 1B-open conformation (Fig. 2).

In three out of the five simulations, the 1B domain underwent a ~90° rotation from the starting 1B-open conformation around the stalk toward the RNA-binding groove to a 1B-closed conformation (Fig. 6A). These ~90° rotated 1B domain conformations closely resemble the disposition of the 1B domain in the nsp13.1-apo structure. The 1B domain rmsd between the simulation-generated structures from the last 2 μs of the three trajectories and the 1B domain in the nsp13.1-apo cryo-EM structure (aligned on the RecA1 domain) was, on average, ∼3.6 Å. We also observed that a small portion (<5%) of these 1B-closed structures were more similar to the 1B domain of the nsp13.1-engaged conformation (rmsd <3.5 Å). These nsp13.1-engaged-like 1B conformations were short-lived, and once visited they quickly transitioned to the nsp13.1-apo conformation, presumably because the nsp13.1-engaged conformation was captured in the presence of RNA, which was not included in our simulations.

**Fig. 6.**
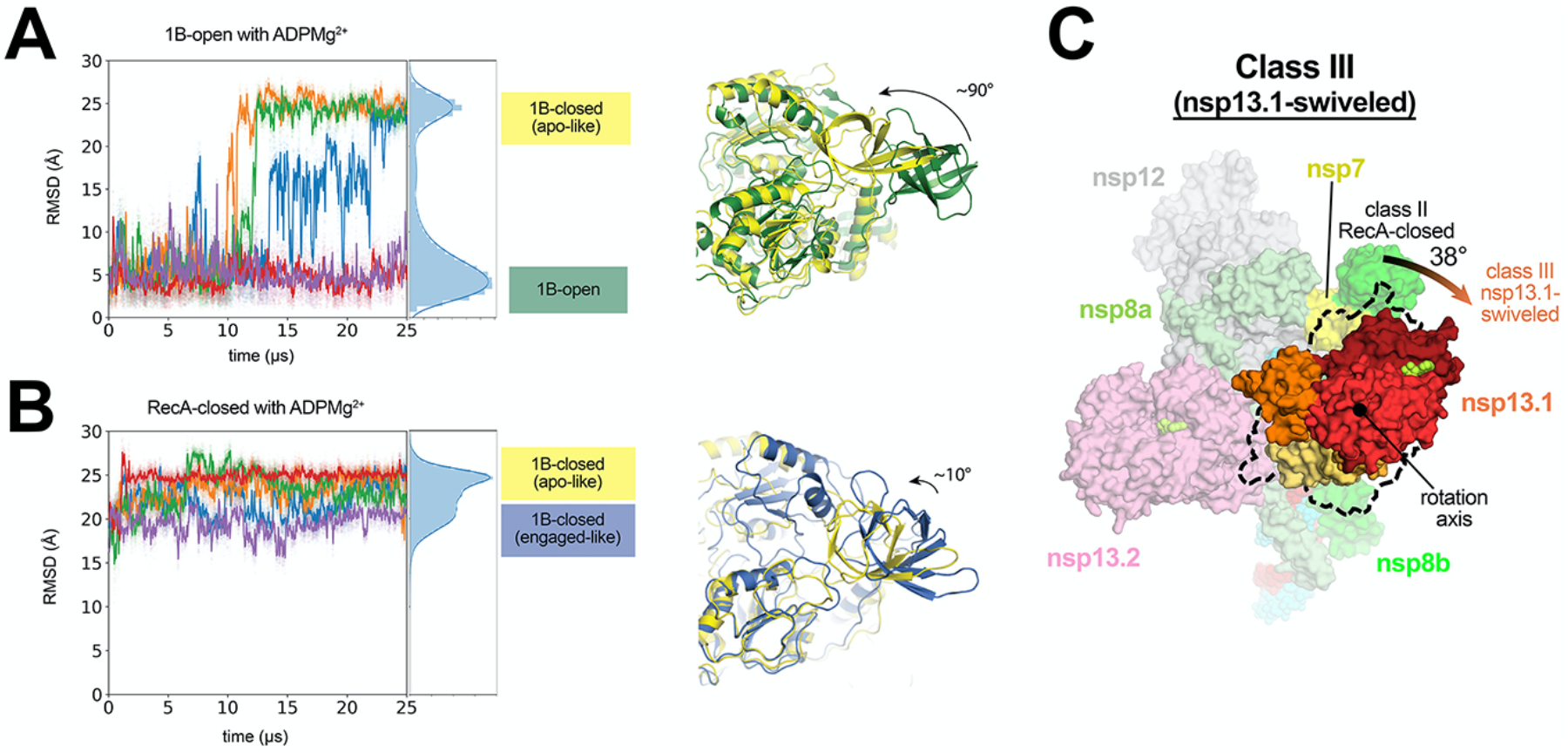
In the nsp13.1-swiveled structure, the entire nsp13.1 promoter is rotated. **A.** Front view of the nsp13.1-swiveled structure, highlighting nsp13.1. The position of the nsp13.1 promoter in the nsp13.1-engaged structure is illustrated by the dashed black outline. The nsp13.1 promoter of the nsp13.1-swiveled structure is rotated by 38° as shown. **B.** Five independent simulations of ADPMg^2+^-bound nsp13.1, starting from the 1B-open cryo-EM structure. Values plotted represent the heavy-atom rmsd of the 1B domain (nsp13 residues 150-228) compared to the 1B domain in the 1B-open cryo-EM structure (aligned on the RecA1 domain). The rmsd histograms on the right represent aggregate values across all five simulations. Representative structures of the two major conformations from the rmsd histogram from simulations are shown (right). **C.** Five independent simulations of ADPMg^2+^-bound nsp13.1, starting from the nsp13.1-engaged state. Values plotted represent the heavy-atom rmsd of the 1B domain compared to the 1B domain in the 1B-open cryo-EM state (aligned on the RecA1 domain). The rmsd histograms on the far right represent aggregate values across all five simulations. Representative structures of the two major conformations from the rmsd histogram from simulations are shown (right). See also Fig. S6.

We next asked whether or not the 1B domain in a 1B-closed state may spontaneously transition to the 1B-open state. In each of the three simulations in which we observed a transition of the 1B domain from the 1B-open to a 1B-closed conformation, the 1B domain remained in the 1B-closed conformation through the end of the 25-μs simulation; a 1B-closed-to-open transition was not observed. We performed an additional five independent 25-μs simulations of the isolated (free) nsp13.1, initiated from the 1B-closed conformation (of the nsp13.1-engaged structure). We did not observe any transition events to the 1B-open conformation over the aggregated 125 μs simulation time. Instead, the 1B domain maintained its 1B-closed orientation in the initial structure, with some minor wobbling back and forth between the 1B-closed conformations of the nsp13.1-engaged and nsp13.1-apo structures (Fig. 6B).

Aligning the nsp13.1 simulation structures in the 1B-open-to-closed transition pathways with the nsp13.1 of the 1B-open cryo-EM structure showed that, on average, ~40% (53%, 22%, and 45% in the three simulations) of these 1B domain intermediate conformations clashed with nsp13.2, suggesting that the 1B-open-to-closed transition might be blocked by nsp13.2 (Fig. S6). Here we envisage that 1B domain transitions are facilitated by entry into the ‘swiveled’ state. The swiveled structure is characterized by one dominant conformational change; compared to the nsp13.1-engaged structure, the nsp13.1 protomer as a whole swivels with respect to the rest of the RTC by 38°, repositioning nsp13.1 with respect to nsp13.2 (Fig. 6C).

There are some clashes between nsp13.1 and nsp13.2 when the simulation-generated structures are aligned to the swiveled cryo-EM structure, but to a much lesser extent (9%, 0%, and 2% in the three simulations; Fig. S6). This observation is consistent with the notion that the swiveled structure may be an intermediate state that facilitates the transition between the 1B-open and 1B-closed conformations.

In summary, our simulations suggest that the conformation of the 1B domain in the 1B-open structure may only be transiently stable on its own, transitioning spontaneously into the 1B-closed conformations of the nsp13.1-apo and nsp13.1-engaged structures. Such transitions may be blocked by the presence of nsp13.2 in the 1B-open nsp13_2_-RTC. We did not observe transitions from the 1B-closed conformations to the 1B-open conformation, and we speculate that in the presence of RNA in the nsp13.1 RNA-binding groove (Fig. 3), nsp13.1 may be further stabilized in the closed 1B domain conformation.

### Nsp13 conformations in nsp13_2_-backtracked complexes

In the nsp13.1-engaged state (Fig. 2), the RdRp translocates in the 3′->5′ direction on the t-RNA while nsp13.1 grasps the single-stranded t-RNA ahead of the RdRp (Fig. 3) and translocates in the 5′->3′ direction (Fig. 1B). We proposed that events at the RdRp active site that would delay or stall p-RNA chain elongation (such as misincorporation or incorporation of nucleotide analogs) could allow the nsp13.1 translocation activity to push the RdRp backward on the t-RNA ^20^. In this process, termed backtracking, the complex moves in the 5′->3′ direction on the t-RNA accompanied by reverse-threading of the p-RNA through the complex, generating a single-stranded p-RNA 3′-fragment. In support of this hypothesis, structural and functional studies showed that the SARS-CoV-2 RdRp can backtrack, that the resulting single-stranded p-RNA 3′-fragment extrudes out the RdRp NTP-entry tunnel, and that backtracking is stimulated by nsp13 ^26^.

To compare the conformational states of the nsp13 protomers in the nsp13_2_-BTCs (backtracked complexes) with the nsp13_2_-RTCs, we used the same masked classification with signal subtraction protocol (Fig. S2) to classify the nsp13_2_-BTC particles into four conformational states (Fig. S7). Structural models were built and rigid-body refined into the cryo-EM densities for each class except for nsp13_2_-BTC-class2 (13% of the particles), which had very poor cryo-EM density for nsp13.1. To compare these structural models with the nsp13_2_-RTC structures, we aligned the models for each nsp13_2_-BTC model with the nsp13.1-engaged state by superimposing α-carbons of nsp12, yielding rmsds < 0.213 Å. We then calculated rmsds for α-carbons of nsp13.1 and nsp13.2. Both nsp13_2_-BTC-class1 and nsp13_2_-BTC-class4 aligned well with the nsp13.1-engaged nsp13_2_-RTC state (Table S4) and both also had strong density for the downstream t-RNA engaged with nsp13.1 (Fig. S7). Therefore, we classify both of these structures as nsp13.1-engaged-BTCs. The nsp13_2_-BTC-class3 structure had an open 1B domain of nsp13.1 and clearly aligned with the 1B-open-RTC structure (Table S4). Thus, in contrast to the nsp13_2_-RTC structures, which were equally divided between the nsp13.1-engaged and 1B-open states (33% each), the nsp13_2_-BTC structures were heavily skewed towards the nsp13.1-engaged state (72%) vs. the 1B-open state (15%; Fig. S7).

## Discussion

In this work, we observed distinct conformational states of the nsp13 protomers within the SARS-CoV-2 nsp13_2_-RTC, providing functional insights into nsp13 and its complex with the RTC (see Video S2). Like other helicases, nsp13 is a molecular motor that translocates along single-stranded nucleic acid, unwinding structural elements in its path (Mickolajczyk et al., 2020). This process is driven by conformational changes within nsp13 resulting from NTP hydrolysis.

The conformational transition from the nsp13.1-engaged to the nsp13.1-apo structures, observed both by our cryo-EM (Fig. 4A) and MD (Figs. 4C, D) analyses, corresponds to an ~21° rotation of the RecA2 domain with respect to RecA1, opening the gap between the two domains (Fig. 4A; Video S1). The nsp13.1-engaged structure is engaged with the substrate RNA and is trapped in an ‘on-pathway’ conformation of the nucleotide hydrolysis cycle by the non-hydrolyzable ATP analog ADP-AlF_3_. While the nsp13.1-apo structure, being devoid of RNA, is not ‘on-pathway’ *per se*, the 21° opening of the RecA2 domain from the nsp13.1-engaged to nsp13.1-apo conformations matches the disposition of the RecA2 domains in other SF1 helicases, such as human Upf1, a structural homolog of nsp13 ^13,23^. The disposition of the RecA domains of Upf1 with ADP-AlF_3_ and RNA substrate [PDB 2XZO] ^33^ matches the nsp13.1-engaged structure. On the other hand, the RecA domains in a structure of Upf1 with ADP (so likely on-pathway) are opened by a 24° rotation about the same axis as the 21° opening of the nsp13.1-apo RecA domains [PDB 2GK6] ^34^. We thus infer that the nsp13.1-apo conformation reports on an on-pathway conformation of the RecA domains, such as in the ADP-Mg^2+^/RNA-bound state of the translocation cycle (Figs. 4C, D). Due to the opening of the nsp13.1 RecA domains, the center-of-gravity of RecA2 shifts roughly parallel with the RNA backbone by 3.4 Å, corresponding to the rise between stacked RNA bases. This observation is suggestive of an ‘inchworm’ model for translocation (Video S1), as proposed for related SF1 helicase translocation on single-stranded nucleic acids ^14,17,35–39^.

Prior structural analysis of the nsp13_2_-RTC identified that the nsp13.1 helicase and the RdRp translocate on the t-RNA with opposing polarities (Chen et al., 2020). In circumstances where RdRp elongation of the p-RNA is hindered (such as in the event of a misincorporation at the p-RNA 3′-end), nsp13.1 translocation activity could backtrack the RdRp ^20^, as shown by follow-up structural and biochemical analyses ^26^. The opposing polarities of the RdRp and nsp13 translocation activities (Fig. 1B) presented a conundrum that was not addressed by these previous studies; how is it possible for the RdRp to rapidly and efficiently synthesize RNA if it is constantly being opposed by nsp13? Moreover, the predominant complex present in the nsp13-RTC samples is the nsp13_2_-RTC complex (Table S1), but only nsp13.1 was seen to engage with the t-RNA; what is the role of nsp13.2, the second copy of nsp13 in the nsp13_2_-RTC? The work herein suggests answers to both questions.

Maximum likelihood classification approaches revealed four distinct conformations of the nsp13 protomers in the nsp13_2_-RTC (Figs. 2, 7; Videos S1, S2). The nsp13.1-engaged state resolves nsp13.1 clamped onto the single-stranded downstream t-RNA, providing an atomic view of nsp13 engaged with the single-stranded RNA (Fig. 3). The single-stranded t-RNA threads through a tunnel formed by a deep groove between the RecA1 and RecA2 domains and further enclosed by the 1B domain (Fig. 5A).

**Fig. 7.**
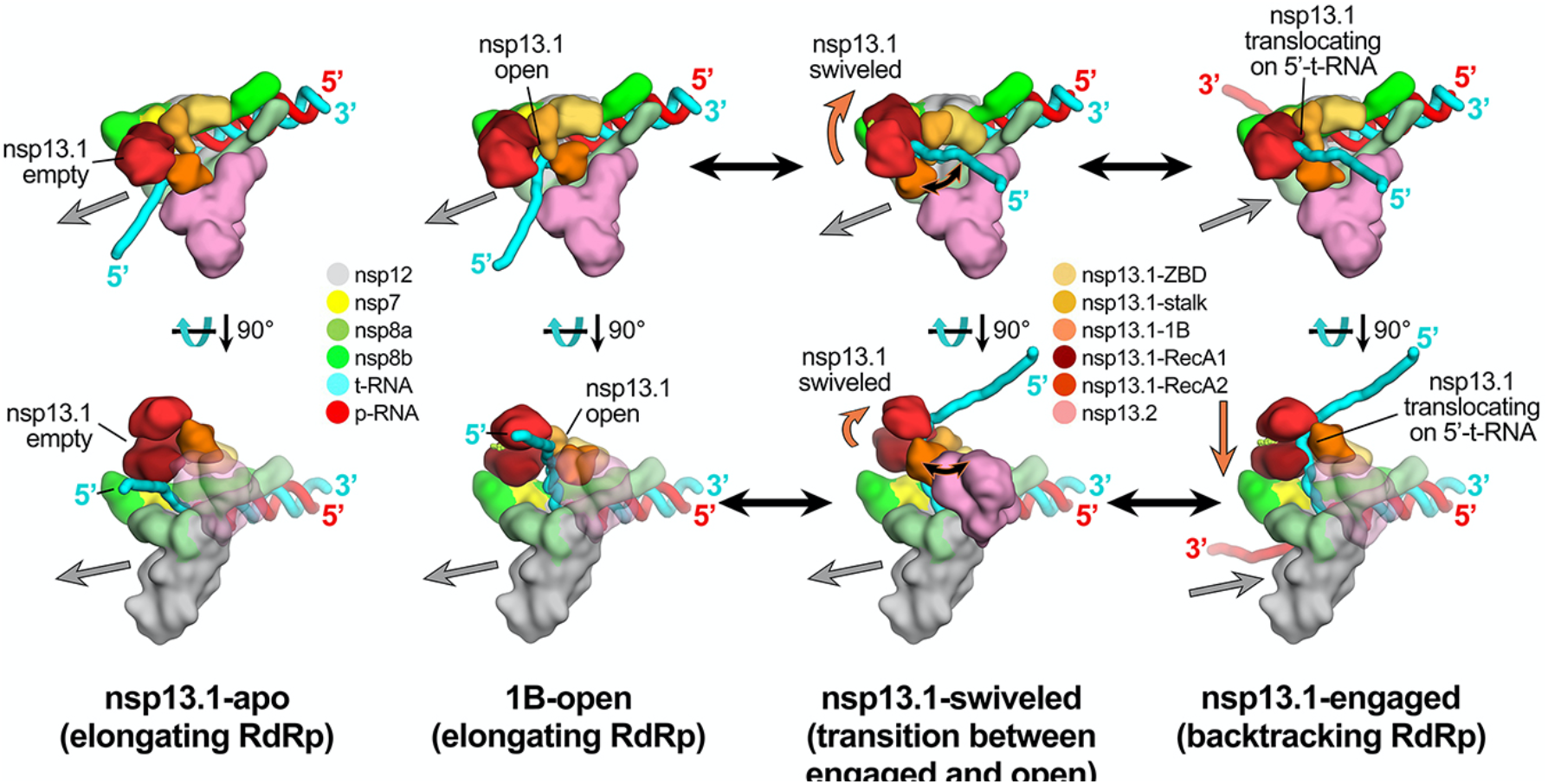
Schematic model for RTC elongation (1B-open) vs. backtracking (nsp13.1-engaged) states. Top views (top row) and side views (bottom row) of each structural class. Nsp13.1-apo (17%): The nsp13.1 RecA domains are open, consistent with the absence of nucleotide. The nsp13.1 is therefore not engaged with the downstream 5′-t-RNA and the RdRp can freely translocate on the t-RNA with concurrent elongation of the p-RNA (gray arrow pointing downstream). 1B-open (33%): The nsp13.1 1B domain is rotated open and sterically trapped by the presence of nsp13.2. The nsp13.1 is therefore unable to engage with the downstream 5′-t-RNA and is inactive. The RdRp is able to elongate freely in the downstream direction. Nsp13.1-swiveled (17%): The rotation of the nsp13.1 protomer away from nsp13.2 provides space for the nsp13.1 1B domain to open and/or close. We therefore propose that nsp13.1-swiveled represents a transition state between the 1B-open (elongating) and nsp13.1-engaged (backtracking) states. Nsp13.1-engaged (33%): The nsp13.1 1B and RecA domains are clamped onto the downstream 5′-t-RNA. In this state, nsp13.1 can translocate on the t-RNA in the 5′-3′ direction (shown by the orange arrow). This counteracts RdRp elongation and causes backtracking (backward motion of the RdRp on the RNA, shown by the gray arrow pointing upstream). Also see Fig. S7 and Videos S1 and S2.

By contrast, the 1B-open state shows nsp13.1 adopting a conformation in which the 1B domain is rotated open ~85° about the stalk domain, leaving an open RNA binding groove (Fig. 5B). In this state, the single-stranded downstream t-RNA does not engage with the helicase. Thus, this represents an inactive state of the helicase that would be unable to translocate on the RNA.

Our structural analysis combined with MD simulations confirmed that the conformation of the nsp13.1 1B domain in the 1B-open structure is not stable on its own but is sterically trapped by the presence of nsp13.2, which blocks the conformational change required for 1B domain closure (Fig. 6A). These results suggest that the 1B-open state represents a rapidly elongating state of the nsp13_2_-RTC, where the downstream single-stranded template RNA feeds into the RdRp active site without engaging with nsp13.1. Nsp13.2 may trap the 1B-open (inactive) state of nsp13.1, allowing RdRp elongation to proceed without opposition from the nsp13.1 helicase (Fig. 7). Finally, swiveling of nsp13.1 in the swiveled state allows space for the 1B-open to 1B-closed transition (Fig. 6C), suggesting that the swiveled state represents a transition state between the open and closed states of the 1B domain (Fig. 7). We note that the presence of nsp13.2 in the nsp13.1-engaged state would also block the 1B-closed to 1B-open transition, suggesting how nsp13.2 can enhance the helicase activity of nsp13.1 ^21^

Thus, our results suggest a mechanism for the nsp13_2_-RTC to turn backtracking on and off; switching between rapid RNA synthesis (1B-open state; elongating RdRp; Fig. 7) and backtracking (nsp13.1-engaged, backtracking RdRp; Fig. 7). In our analysis of the conformational states of the nsp13_2_-RTC, the particles were equally divided between the nsp13.1-engaged (backtracking on) and 1B-open (backtracking off) states (Figs. S2, S7). Remarkably, an identical analysis of the backtracked nsp13_2_-BTC ^26^ revealed a strikingly different distribution of particles in which the nsp13.1-engaged (backtracking on) state was heavily favored (Fig. S7). This raises the possibility that the conformational switch that turns backtracking on and off is allosterically controlled.

In MD simulations exploring the dynamics of the p-RNA 3′-nucleotide of a pre-translocated RTC, a mismatched p-RNA 3′-nucleotide frayed from the t-RNA towards and into the NTP-entry tunnel (which also serves as the backtracking tunnel; Video S1), while a p-RNA 3′-nucleotide engaged in a Watson-Crick base pair with the t-RNA did not ^26^. We thus suggest that misincorporation by the RdRp leads to fraying of the p-RNA 3′-nucleotide into the NTP-entry tunnel, which may allosterically signal the rapidly elongating 1B-open state to switch to the backtracking nsp13.1-engaged state (Fig. 7). This facilitates a possible proofreading mechanism since backtracking would extrude the mismatched p-RNA 3′-nucleotide out of the NTP-entry tunnel (Fig. 7) ^26^, allowing the nsp10/nsp14 3′-exonuclease proofreading activity to access and degrade the mismatched p-RNA 3′-nucleotide ^40–42^. The nsp14-mediated proofreading activity is crucial for the virus to avoid mutation catastrophe while replicating its ~30 kb genome ^41^, and is also an important determinant of SARS-CoV-2 susceptibility to many anti-viral nucleotide analogs ^43^.

## Acknowledgments

We thank M. Ebrahim and L. Urnavicius at The Rockefeller University Evelyn Gruss Lipper Cryo-electron Microscopy Resource Center and H. Kuang at the New York Structural Biology Center (NYSBC) for help with cryo-EM data collection, and R. Landick, T. Appleby, and members of the Darst/Campbell laboratory for helpful discussions, and M. Grasso, P.M.M. Shelton, T. M. Kapoor, P.D.B. Olinares, and B.T. Chait (The Rockefeller University) for helpful discussions and initial sample characterizations and analyses. Some of the work reported here was conducted at the Simons Electron Microscopy Center (SEMC) and the National Resource for Automated Molecular Microscopy (NRAMM) and National Center for CryoEM Access and Training (NCCAT) located at the NYSBC, supported by grants from the NIH National Institute of General Medical Sciences (P41 GM103310), NYSTAR, the Simons Foundation (SF349247), the NIH Common Fund Transformative High Resolution Cryo-Electron Microscopy program (U24 GM129539) and NY State Assembly Majority. This work was supported by the Pels Family Center for Biochemistry and Structural Biology (The Rockefeller University), and NIH grants R01 GM114450 (E.A.C.), R35 GM118130 (S.A.D.), and R01 AI161278 (E.A.C./S.A.D).

## Author contributions

Conceptualization; J.C., Q.W., B.M., J.P., E.A.C., D.E.S., S.A.D.; Cloning, protein purification, biochemistry; J.C., B. M., E.L.; Cryo-EM specimen preparation; J.C., B.M., E.L.; Cryo-EM data collection and processing: J.C., K.M., E.T.E.; Model building and structural analysis: J.C., B.M., E.A.C., S.A.D.; Molecular dynamics simulation and analysis; Q.W., Y.P.; Funding acquisition and supervision: E.A.C., D.E.S., S.A.D.’ Manuscript first draft: Q.W., B.M., J.P., E.A.C., D.E.S., S.A.D.; All authors contributed to finalizing the written manuscript.

## METHODS

No statistical methods were used to predetermine sample size. The experiments were not randomized, and the investigators were not blinded to allocation during experiments and outcome assessment.

### Protein expression and purification

*SARS-CoV-2 nsp12* was expressed and purified as described ^20^. Briefly, a pRSFDuet-1 plasmid containing His_6_-SUMO SARS-CoV-2 nsp12 (Addgene #159107) was transformed into *E. coli* BL21-CodonPlus cells (Agilent). Cells were grown and protein expression was induced by the addition of isopropyl β-d-1-thiogalactopyranoside (IPTG). Cells were collected and lysed in a French press (Avestin). The lysate was cleared by centrifugation and purified on a HiTrap Heparin HP column (Cytiva). The fractions containing nsp12 were loaded onto a HisTrap HP column (Cytiva) for further purification. Eluted nsp12 was dialyzed, cleaved with His_6_-Ulp1 SUMO protease, and passed through a HisTrap HP column to remove the SUMO protease. Flow-through was collected, concentrated by centrifugal filtration (Amicon), and loaded on a Superdex 200 Hiload 16/600 (Cytiva). Glycerol was added to the purified nsp12, aliquoted, flash-frozen with liquid N_2_, and stored at −80°C.

*SARS-CoV-2 nsp7/8* was expressed and purified as described ^20^. Briefly, the pCDFDuet-1 plasmid containing His_6_ SARS-CoV-2 nsp7/8 (Addgene #159092) was transformed into *E. coli* BL21 (DE3). Cells were grown and protein expression was induced by the addition of IPTG. Cells were collected and lysed in a French press (Avestin). The lysate was cleared by centrifugation and purified on a HisTrap HP column (Cytiva). Eluted nsp7/8 was dialyzed, cleaved with His_6_-Prescission Protease to cleave His_6_ tag, and then passed through a HisTrap HP column to remove the protease (Cytiva). Flow-through was collected, concentrated by centrifugal filtration (Amicon), and loaded onto a Superdex 75 Hiload 16/600 (Cytiva). Glycerol was added to the purified nsp7/8, aliquoted, flash-frozen with liquid N_2_, and stored at −80°C.

*SARS-CoV-2 nsp13* was expressed and purified as described ^20^. Briefly, the pet28 plasmid containing His_6_ SARS-CoV-2 nsp13 (Addgene #159390) was transformed into *E. coli* Rosetta (DE3) (Novagen). Cells were grown and protein expression was induced by the addition of IPTG. Cells were collected and lysed in a French press (Avestin). The lysate was cleared by centrifugation and purified on a HisTrap HP column (Cytiva). Eluted nsp13 was dialyzed, cleaved with His_6_-Prescission Protease, and then passed through a HisTrap HP column to remove protease (Cytiva). Flow-through was collected, concentrated by centrifugal filtration (Amicon), and loaded onto a Superdex 200 Hiload 16/600 (Cytiva). Glycerol was added to the purified nsp13, aliquoted, flash-frozen with liquid N_2_, and stored at −80°C.

#### Preparation of SARS-CoV-2 nsp13-replication/transcription complex (RTC) for Cryo-EM

Cryo-EM samples of SARS-CoV-2 nsp13-RTC were prepared as described ^20^. Briefly, purified nsp12 and nsp7/8 were concentrated, mixed in a 1:3 molar ratio, and incubated for 20 min at 22°C. Annealed RNA scaffold (Horizon Discovery, Ltd.) was added to the nsp7/8/12 mixture and incubated for 15 min at 22°C. Sample was buffer exchanged into cryo-EM buffer [20 mM HEPES pH 8.0, 150 mM K-Acetate,10 mM MgCl_2_, 2 mM DTT] and further incubated for 20 min at 30°C. The sample was purified over a Superose 6 Increase 10/300 GL column (Cyriva) in cryo-EM buffer. The peak corresponding to nsp7/8/12/RNA complex was pooled and concentrated by centrifugal filtration (Amicon). Purified nsp13 was concentrated by centrifugal filtration (Amicon) and buffer exchanged into cryo-EM buffer. Buffer exchanged nsp13 was mixed with ADP (1 mM final) and AlF_3_ (1 mM final) and then added to nsp7/8/12/RNA at a molar ratio of 1:1. Complex was then incubated for 5 min at 30°C.

#### Cryo-EM grid preparation

Prior to grid freezing, 3-([3-cholamidopropyl]dimethylammonio)-2-hydroxy-1-propanesulfonate (CHAPSO, Anatrace) was added to the sample (8 mM final), resulting in a final complex concentration of 8 μM. The final buffer condition for the cryo-EM sample was 20 mM HEPES pH 8.0, 150 mM K-Acetate,10 mM MgCl_2_, 2 mM DTT, 1 mM ADP, 1 mM AlF_3_, 8 mM CHAPSO. C-flat holey carbon grids (CF-1.2/1.3-4Au, EMS) were glow-discharged for 20 s prior to the application of 3.5 μL of sample. Using a Vitrobot Mark IV (Thermo Fisher Scientific), grids were blotted and plunge-froze into liquid ethane with 90% chamber humidity at 4°C.

#### Cryo-EM data acquisition and processing

Structural biology software was accessed through the SBGrid consortium ^44^. Grids were imaged using a 300 kV Titan Krios (Thermo Fisher Scientific) equipped with a GIF BioQuantum and K3 camera (Gatan). Images were recorded with Leginon ^45^ with a pixel size of 1.07 Å/px (micrograph dimension of 5760 × 4092 px) over a defocus range of −0.8 μm to −2.5 μm with a 20 eV slit. Movies were recorded in “counting mode” (native K3 camera binning 2) with ∼30 e-/px/s in dose-fractionation mode with subframes of 50 ms over a 2.5 s exposure (50 frames) to give a total dose of ∼66 e-/Å^2^. Dose-fractionated movies were gain-normalized, drift-corrected, summed, and dose-weighted using MotionCor2 ^46^. The contrast transfer function (CTF) was estimated for each summed image using the Patch CTF module in cryoSPARC v2.15.0 ^47^. Particles were picked and extracted from the dose-weighted images with box size of 256 px using cryoSPARC Blob Picker and Particle Extraction. The entire dataset consisted of 17,806 motion-corrected images with 3,750,107 particles. Particles were sorted using two rounds of cryoSPARC 2D classification (N=100, where N equals the number of classes), resulting in 661,105 curated particles that were re-extracted with a boxsize of 320 px. An initial model was generated using cryoSPARC *Ab initio* Reconstruction (N=3) on a subset of the particles. Particles were further curated using this initial model as a 3D template for cryoSPARC Heterogeneous Refinement (N=3), resulting in 451,760 particles (green map, Fig. S1). Curated particles were further classified using cryoSPARC Heterogeneous Refinement (N=3). Each of the resulting 3D classes were further processed with cryoSPARC *Ab initio* Reconstruction (N=3), generating three distinct models that could be used to sort particles [Ref 1: nsp13_1_-RTC, Ref 2: nsp13_2_-RTC, Ref 3: (nsp13_2_-RTC)_2_]. Using Ref 1-3 as 3D templates for Heterogeneous Refinement (N=6), multi-reference classification was performed on the 451,760 curated particles. Classification revealed three unique classes: nsp13_1_-RTC (class1; 85,206 particles; yellow), nsp13_2_-RTC (class2-4; 315,216 particles; red), and (nsp13_2_-RTC)_2_ (class5; 35,403 particles; blue). Particles within each class were further processed using RELION 3.1-beta Bayesian Polishing ^48^. Polished particles were refined using cryoSAPRC Local and Global CTF Refinement in combination with cryoSPARC Non-uniform Refinement 49, resulting in structures with the following particle counts and nominal resolutions: nsp13_1_-RTC (85,187 particles; 3.2 Å), nsp13_2_-RTC (315,120 particles; 2.9 Å), (nsp13_2_-RTC)_2_ (35,392 particles; 3.3 Å). To facilitate model building of nsp13_2_-RTC, particles from nsp13_1_-RTC and nsp13_2_-RTC were combined in a cryoSPARC Non-uniform Refinement, subtracted (masking the RTC), and further refined with cryoSPARC Local Refinement using a mask encompassing the RTC. The resulting map, deemed RTC (local), had nominal resolution of 2.8 Å. Additionally, particles from the nsp13_2_-RTC were subtracted in different regions (using separate masks for nsp12-NiRAN, nsp13.1, and nsp13.2) and the particles from each subtraction were further refined with masked cryoSPARC Local Refinement. The resulting maps had the following nominal resolutions: nsp13.1(local): 3.4 Å, nsp13.2(local): 3.3 Å, nsp12-NiRAN(local): 2.7 Å. Locally refined maps were combined into an nsp13_2_-RTC composite map using PHENIX ‘Combine Focused Maps’ ^50,51^, with resulting nominal resolution of 2.8 Å. The nsp13-RecA domains in particles from the nsp13_1_-RTC and nsp13_2_-RTC classes were sorted using particle subtraction (masking around the RecA domains, shown as red mesh in Fig. S1), followed by masked RELION 3D classification. Classification of RecA domains in the nsp13_1_-RTC particles (pale yellow) did not reveal discrete conformational heterogeneity in the RecA domains. However, classification of RecA domains in the nsp13_2_-RTC particles (light red) revealed unique conformations of the RecA domains with the following particle counts and nominal resolutions: RecA classI (52,403 particles; 3.5 Å), RecA class II (102,615 particles; 3.1 Å), RecA class III (54,830 particles; 3.5 Å), RecA class IV (105,272 particles; 3.1 Å). Local resolution calculations were generated using blocres and blocfilt from the Bsoft package ^29^.

#### Model building and refinement

For an initial model of the nsp13_2_-RTC, the initial RTC model was derived from PDB 6XEZ ^20^ and the initial nsp13 model from PDB 6ZSL ^32^. The models were manually fit into the cryo-EM density maps using Chimera ^52^ and rigid-body and real-space refined using Phenix real-space-refine ^50,51^. For real-space refinement, rigid body refinement was followed by all-atom and B-factor refinement with Ramachandran and secondary structure restraints. Models were inspected and modified in Coot ^53^.

### Molecular dynamics simulations

#### General simulation setup and parameterization

Proteins, ATP, ADP, and ions were parameterized with the DES-Amber SF1.0 force field ^54^. RNAs were parameterized with the Amber ff14 RNA force field ^55^ with modified electrostatic, van der Waals, and torsional parameters to more accurately reproduce the energetics of nucleobase stacking ^56^. The systems were solvated with water parameterized with the TIP4P-D water model ^57^ and neutralized with a 150 mM NaCl buffer. The systems each contained ~160,000 atoms in a 110 × 110 × 110 Å cubic box.

Systems were first equilibrated on GPU Desmond using a mixed NVT/NPT schedule ^58^, followed by a 1 μs relaxation simulation on Anton, a special-purpose machine for molecular dynamics simulations ^59^. All production simulations were performed on Anton and initiated from the last frame of the relaxation simulation. Production simulations were performed in the NPT ensemble ^60^ at 310 K using the Martyna-Tobias-Klein barostat ^61^. The simulation time step was 2.5 fs, and a modified r-RESPA integrator ^62^ was used in which long-range electrostatic interactions were evaluated every three time steps. Electrostatic forces were calculated using the *u*-series method ^63^. A 9-Å cutoff was applied for the van der Waals calculations.

#### System preparation

The initial conformations of Class II nsp13.1 bound to the various substrates (ATPMg^2+^/RNA, ADPMg^2+^/RNA, ATPMg^2+^, and ADPMg^2+^) were prepared based on the cryo-EM structure of the Class II nsp13_2_-BTC_5._ The initial conformation of the Class IV, 1B-open nsp13.1 structure was prepared from the cryo-EM Class IV nsp13_2_-BTC_5_ structure. AlF_3_ was removed from the active site. Missing loops and termini in proteins were capped with ACE/NME capping groups. In simulations with ATP at the active site, ATP was manually placed using ADP in the cryo-EM structure as the reference. The systems were prepared for simulation using the Protein Preparation Wizard in Schrödinger Maestro (Schrödinger Release 2020-4: Maestro, Schrödinger, LLC, New York, NY, 2020).

#### Simulation analysis

The average rmsd was calculated for the RecA2 domain (residues 450–690) and 1B domain (residues 145–200) of nsp13.1 between the cryo-EM structures and instantaneous structures from the trajectories, aligned on the RecA1 lobe (residues 240–440). Simulation structures shown in figures were rendered using PyMol (The PyMOL Molecular Graphics System, Version 2.0 Schrödinger, LLC).

### Quantification and statistical analysis

The local resolution of the cryo-EM maps (Figs S3 and S5) was estimated using blocres ^29^ with the following parameters: box size 15, sampling 1.1, and cutoff 0.5. Directional 3DFSCs (Figs. S3 and S5) were calculated using 3DFSC ^64^. The quantification and statistical analyses for model refinement and validation were generated using MolProbity ^65^ and PHENIX ^51^.

### Data and code availability

All unique/stable reagents generated in this study are available without restriction from one of the Lead Contacts, Seth A. Darst. The cryo-EM density maps and atomic coordinates have been deposited in the EMDataBank and Protein Data Bank as follows: nsp13_1_-RTC (EMD-24431, 7RE2), nsp13_2_-RTC (composite) (EMD-24430, 7RE1), (nsp13_2_-RTC)_2_ (EMD-24432, 7RE3), nsp13_2_-RTC (nsp13.1-apo) (EMD-24428, 7RDZ), nsp13_2_-RTC (nsp13.1-engaged) (EMD-24427, 7RDY), nsp13_2_-RTC (nsp13.1-swiveled) (EMD-24429, 7RE0), nsp13_2_-RTC (1B-open) (EMD-24426, 7RDX). The MD trajectories described in this work are available at https://www.deshawresearch.com/downloads/download_trajectory_sarscov2.cgi/.

