## Supplement for "Ensemble cryo-electron microscopy reveals conformational states of the nsp13 helicase in the SARS-CoV-2 helicase replication-transcription complex"

Supplemental information includes 4 tables, 7 figures, and 2 videos.

**Table S1. Cryo-EM datasets of the nsp13-RTC. Related to Fig. 1.**

| Structural class | Dataset 1 [1] | | | Dataset 2 | | |
|---|---|---|---|---|---|---|
| | particles (%) | nominal resolution (Å)[1] | | particles (%) | map | nominal resolution (Å)[1] |
| nsp13$_1$-RTC | 17,345 (20%) | 4.0 | | 85,187 (20%) | 2 | 3.2 |
| nsp13$_2$-RTC | 58,942 (67%) | 3.5 | | 315,120 (72%) | 3 | 2.9 |
| (nsp13$_2$-RTC)$_2$ | 11,771 (13%) | 7.9 | | 35,392 (8%) | 4 | 3.3 |
| total particles | 88,058 | | | 435,699 | | |

[1]Gold-standard FSC calculated by RELION [2].

**Table S2 | Cryo-EM data collection, refinement and validation statistics for $nsp13_1$-RTC, $nsp13_2$-RTC, and $(nsp13_2\text{-RTC})_2$. Related to Fig. 1.**

| **Dataset** | | **nsp13-RTC** | |
|---|---|---|---|
| **Sample ID** | $nsp13_1$-RTC | $nsp13_2$-RTC (composite) | $(nsp13_2\text{-RTC})_2$ |
| **EMDB** | EMD-24431 | EMD-24430 | EMD-24432 |
| **PDB** | 7RE2 | 7RE1 | 7RE3 |
| **Data collection and processing** | | | |
| Microscope | | FEI Titan Krios | |
| Voltage (kV) | | 300 | |
| Detector | | Gatan K3 | |
| Electron exposure (e–/Å2) | | 66 | |
| Defocus range (µm) | | -0.8 to -2.5 | |
| Data collection mode | | Counting Mode | |
| Nominal Magnification | | 81,000x | |
| Pixel size (Å) | | 1.07 | |
| Symmetry imposed | | C1 | |
| Initial particle images (no.) | | 3,750,107 | |
| Final particle images (no.) | 85,187 | 315,120 | 35,392 |
| Map resolution (Å) - FSC threshold 0.143 | 3.2 | 2.8 | 3.3 |
| Map resolution range (Å) | 2.7-6.2 | 2.1-6.7 | 2.9-6.4 |
| **Refinement** | | | |
| Initial models used (PDB code) | 6XEZ, 6YYT, 6ZSL | 6XEZ, 6YYT, 6ZSL | 6XEZ, 6YYT, 6ZSL |
| Map sharpening B factor (Å2) | -81.9 | -61.1 | -97.4 |
| Model composition | | | |
| Non-hydrogen atoms | 17,096 | 21,825 | 43,154 |
| Protein residues | 1,963 | 2,553 | 5,106 |
| Nucleic acid residues (RNA) | 71 | 71 | 142 |
| Ligands | 5 $Zn^{2+}$, 2 $Mg^{2+}$, 3 CHAPSO, 2 ADP, $AlF_3$ | 8 $Zn^{2+}$, 3 $Mg^{2+}$, 3 CHAPSO, 3 ADP, 2 $AlF_3$ | 16 $Zn^{2+}$, 6 $Mg^{2+}$, 2 CHAPSO, 6 ADP, 4 $AlF_3$ |
| B factors ($Å^2$) | | | |
| Protein | 67.76 | 88.73 | 151.00 |
| Nucleic acid | 151.77 | 175.13 | 158.63 |

| | | | |
|---|---|---|---|
| Ligands | 76.87 | 87.6 | 200.51 |
| R.m.s. deviations | | | |
| Bond lengths (Å) | 0.005 | 0.004 | 0.004 |
| Bond angles (°) | 0.591 | 0.655 | 0.997 |
| Validation | | | |
| Clashscore | 8.79 | 7.84 | 10.86 |
| Poor rotamers (%) | 2.42 | 8.64 | 6.67 |
| Ramachandran plot | | | |
| Favored (%) | 94.98 | 94.02 | 93.68 |
| Allowed (%) | 5.02 | 5.98 | 6.32 |
| Disallowed (%) | 0.0 | 0.0 | 0.0 |

**Table S3 | Cryo-EM data collection, refinement and validation statistics for nsp13$_1$-RTC, nsp13$_2$-RTC, and (nsp13$_2$-RTC)$_2$. Related to Fig. 2.**

| **Dataset** | | **nsp13$_2$-RTC** | | |
|---|---|---|---|---|
| Initial particle images (no.) | | 315,120 | | |
| sample ID | nsp13.1-apo | nsp13.1-engaged | nsp13.1-swiveled | 1B-open |
| EMDB | EMD-24428 | EMD-24427 | EMD-24429 | EMD-24426 |
| PDB | 7RDZ | 7RDY | 7RE0 | 7RDX |
| Final particle images (no.) | 52,403 | 102,615 | 54,830 | 105,272 |
| Map resolution (Å) - FSC threshold 0.143 | 3.6 | 3.1 | 3.5 | 3.1 |
| Map resolution range (Å) | 3.0-7.2 | 2.6-6.9 | 3.0-6.6 | 2.8-6.2 |
| **Refinement** | rigid body | all atom | rigid body | all atom |
| Initial model used (PDB code) | 6XEZ, 6ZSL | 6XEZ, 6ZSL | 6XEZ, 6ZSL | 6XEZ, 6ZSL |
| Map sharpening B factor (Å$^2$) | -78.3 | -84.3 | -75.8 | -82.3 |
| Model composition | | | | |
| Non-hydrogen atoms | 21,545 | 21,846 | 21,563 | 21,828 |
| Protein residues | 2,553 | 2,553 | 2,553 | 2,553 |
| Nucleic acid residues (RNA) | 71 | 78 | 71 | 80 |
| Ligands | 8 $Zn^{2+}$, 2 $Mg^{2+}$,<br>2 ADP, $AlF_3$ | 8 $Zn^{2+}$, 3 $Mg^{2+}$,<br>3 CHAPSO,<br>3 ADP, 2 $AlF_3$ | 8 $Zn^{2+}$, 3 $Mg^{2+}$,<br>3 ADP, 2 $AlF_3$ | 8 $Zn^{2+}$, 3 $Mg^{2+}$,<br>3 CHAPSO,<br>3 ADP, 2 $AlF_3$ |
| B factors (Å$^2$) | | | | |
| Protein | - | 99.83 | - | 129.95 |
| Nucleic acid | - | 144.22 | - | 165.04 |
| Ligands | - | 97.16 | - | 129.50 |
| R.m.s. deviations | | | | |

| | | | | |
|---|---|---|---|---|
| Bond lengths (Å) | 0.015 | 0.015 | 0.041 | 0.017 |
| Bond angles (°) | 0.981 | 0.967 | 1.127 | 1.086 |
| Validation | | | | |
| Clashscore | 11.62 | 8.05 | 12.57 | 8.84 |
| Poor rotamers (%) | 9.10 | 10.18 | 9.10 | 14.58 |
| Ramachandran plot | | | | |
| Favored (%) | 92.94 | 92.40 | 93.01 | 91.64 |
| Allowed (%) | 6.78 | 7.60 | 6.80 | 8.36 |
| Disallowed (%) | 0.28 | 0.0 | 0.20 | 0.0 |

**Table S4. Comparison of $nsp13_2$-BTC and $nsp13_2$-RTC classes.**

| | | | | rmsd (Å) against $nsp13_2$-RTC class | | | |
|---|---|---|---|---|---|---|---|
| BTC-class[a] | particles | resolution (Å) | rms_cur[b] | nsp13.1-apo | nsp13.1-engaged | swiveled | 1B-open |
| class1 | 22,132 (9.4%) | 4.1 | all | 5.955 | 3.715 | 9.071 | 8.372 |
| | | | nsp13.1+nsp13.2 | 8.748 | 5.421 | 13.349 | 12.306 |
| | | | nsp13.1 | 7.8 | 1.493 | 13.282 | 10.941 |
| class2 | 31,364 (13%) | 4.0 | | | | | |
| class3 | 35,082 (15%) | 3.9 | all | 6.689 | 5.801 | 8.325 | 3.156 |
| | | | nsp13.1+nsp13.2 | 9.888 | 8.551 | 12.307 | 4.614 |
| | | | nsp13.1 | 13.211 | 11.201 | 16.101 | 1.716 |
| class4 | 146,569 (62%) | 3.6 | all | 4.626 | 0.685 | 7.099 | 6.246 |
| | | | nsp13.1+nsp13.2 | 6.784 | 0.963 | 10.466 | 9.167 |
| | | | nsp13.1 | 8.776 | 0.303 | 13.057 | 10.854 |

[a]Fig. S7.

[b]After superimposition via the $\alpha$-carbons of nsp12, the $nsp13_2$-BTC components listed in this column were compared with the same components of the $nsp13_2$-RTC class using the PyMOL rms_cur command.

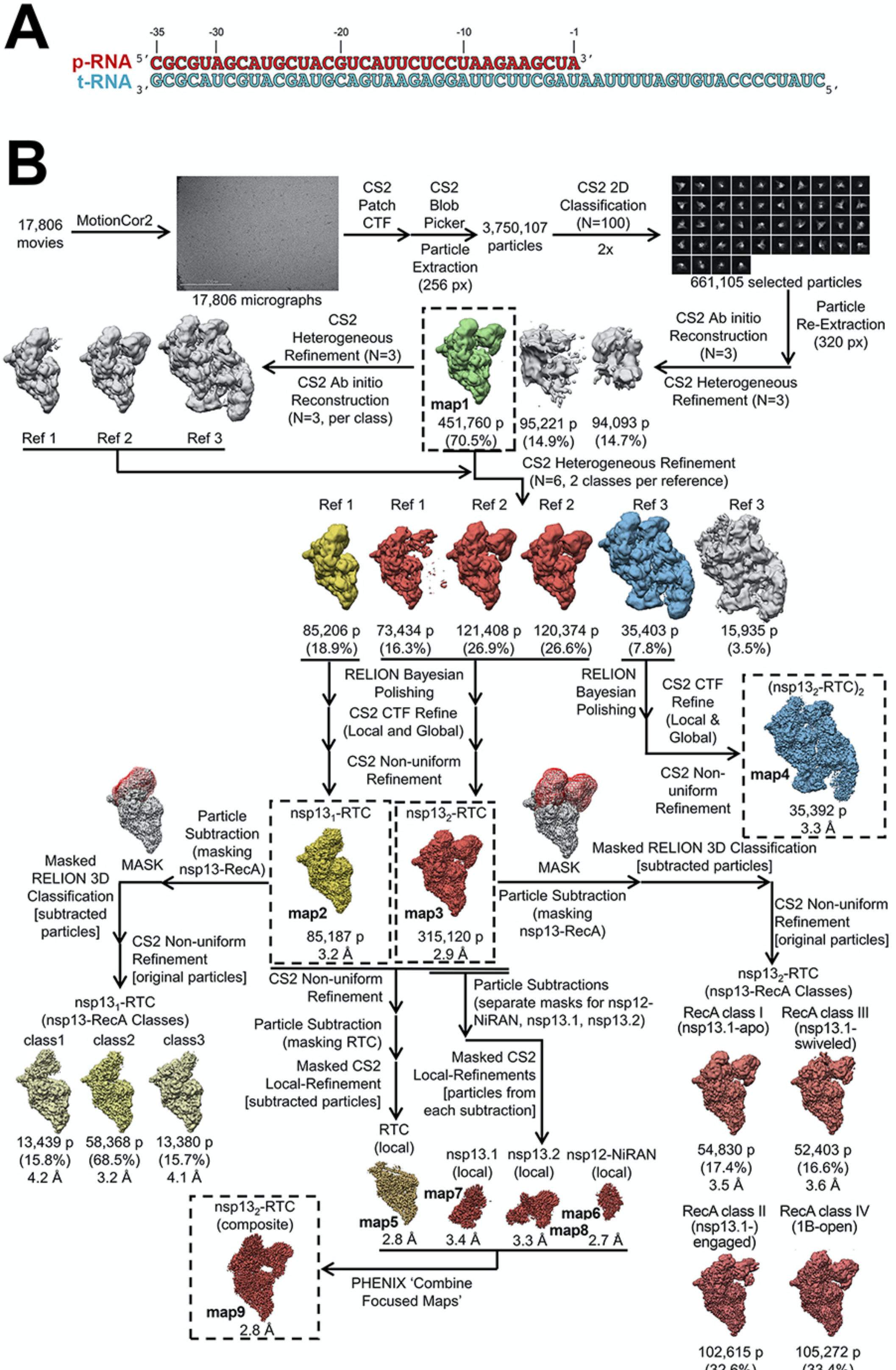

Chen et al., Figure S1

**Fig. S1 | RTC-scaffold and cryo-EM processing pipeline for nsp13-RTC. Related to Fig. 1.**

**A.** RTC scaffold used for RTC cryo-EM.

**B**. Cryo-EM processing pipeline.

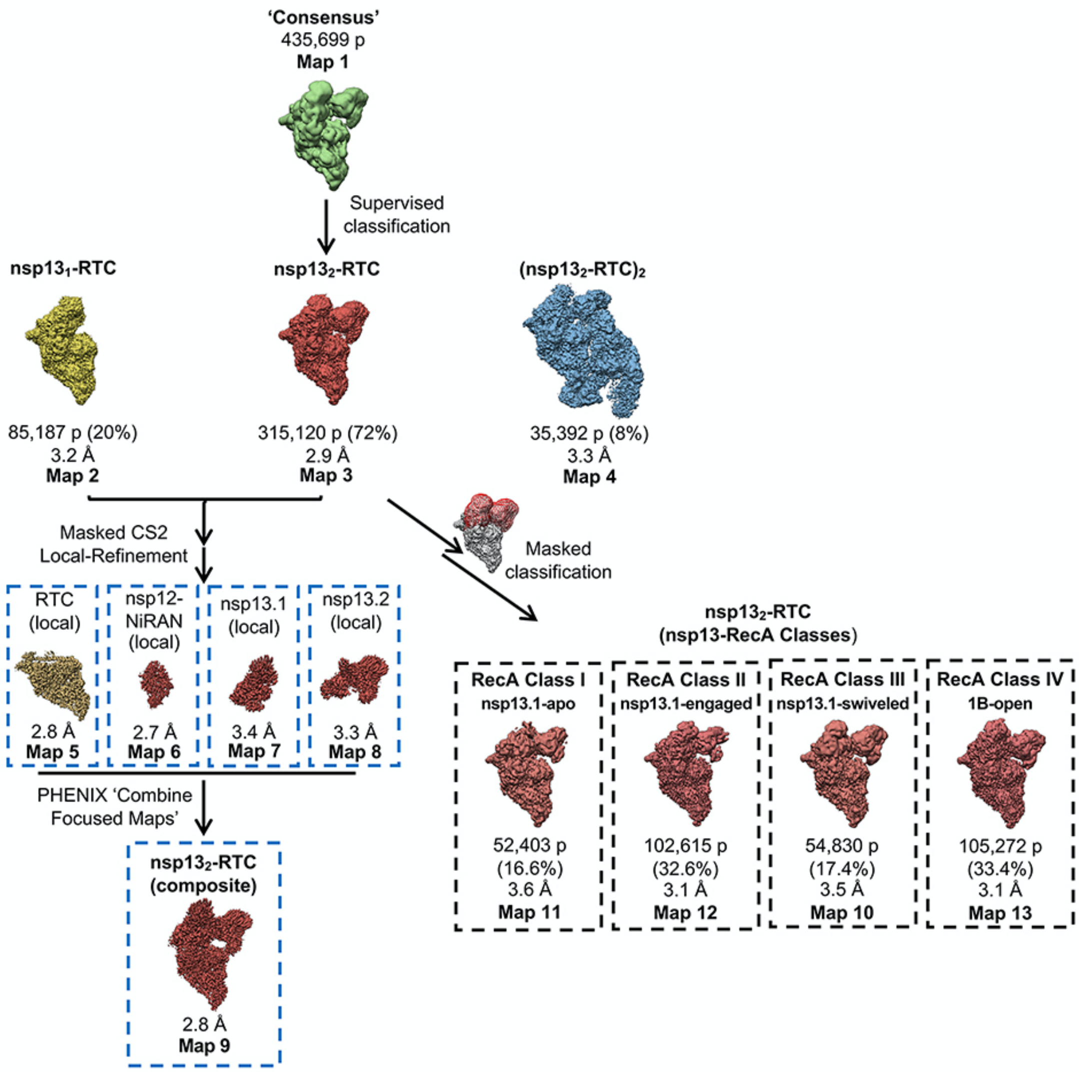

Chen et al., Figure S2

**Fig. S2 | Processing flowchart overview and map nomenclature. Related to Fig. 1.**

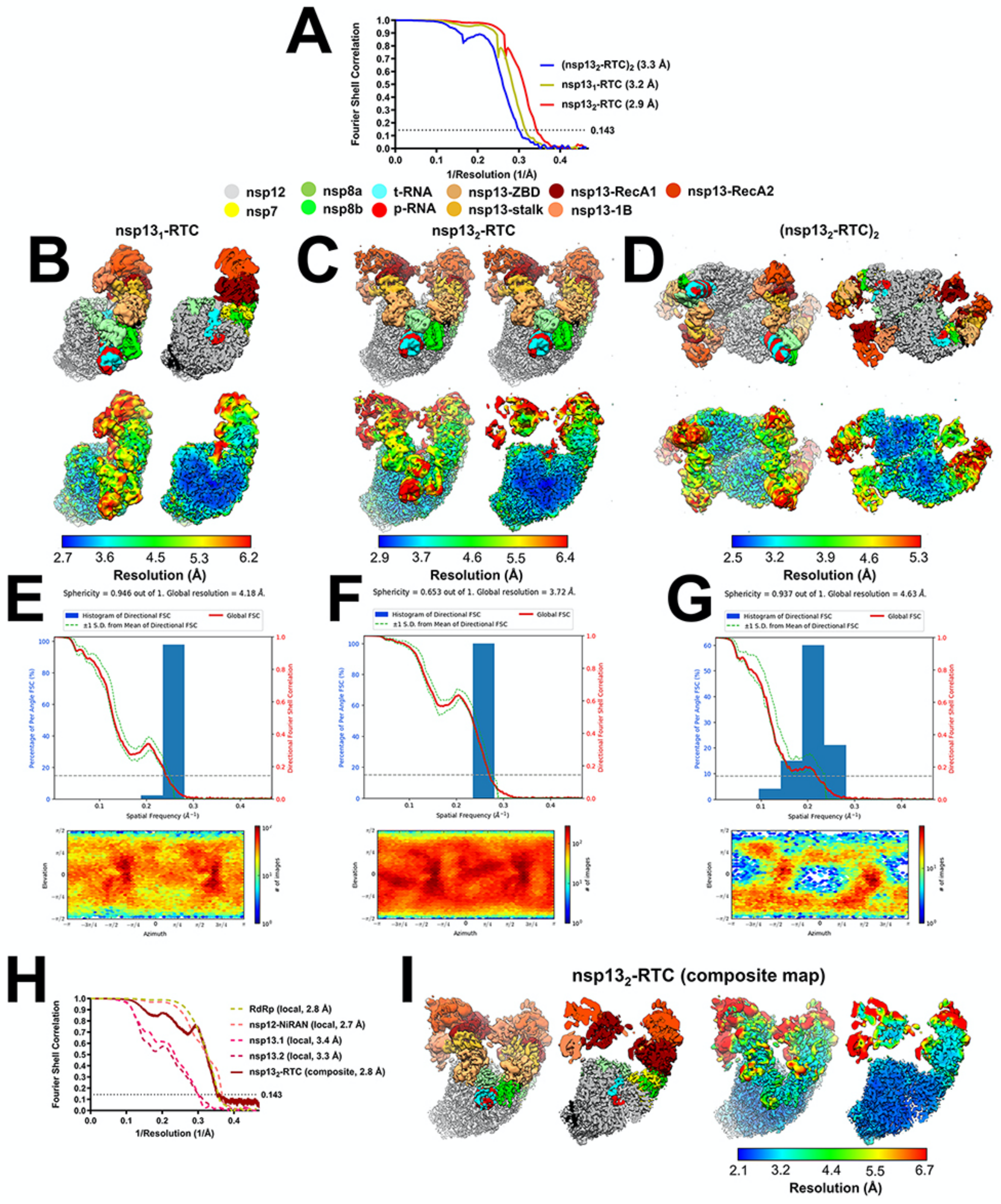

Chen et al., Figure S3

**Fig. S3 | Cryo-EM of nsp13-RTC. Related to Fig. 1.**

**A.** Gold-standard Fourier-shell correlation (FSC) plot for $nsp13_1$-RTC (yellow line), $nsp13_2$-RTC (red line), and ($nsp13_2$-RTC$)_2$ (blue line) calculated by comparing the independently determined half-maps from cryoSPARC. The dotted line represents the 0.143 FSC cutoff which indicates a nominal resolution of 3.2 Å ($nsp13_2$-RTC), 2.9 Å ($nsp13_2$-RTC), and 3.3 Å [($nsp13_2$-RTC$)_2$].

**B - D.** The cryo-EM maps filtered by local resolution [3] are shown. The view on the right is a cross-section.

*(top)* Colored by subunit (key above).

*(bottom)* Colored by local resolution (key on the bottom).

**B.** $Nsp13_1$-RTC (map2).

**C.** $Nsp13_2$-RTC (map3).

**D.** ($nsp13_2$-RTC$)_2$ (map4).

**E - G.** Directional 3D Fourier shell correlation calculated by 3DFSC (*top*) [4] and particle orientation distribution calculated by cryoSPARC (*bottom*).

**E.** $Nsp13_1$-RTC.

**F.** $Nsp13_2$-RTC.

**G.** ($nsp13_2$-RTC$)_2$.

**H.** Gold-standard FSC plots for local-refined maps (dashed lines) and $nsp13_2$-RTC composite map (map9) (solid line).

**I.** The $nsp13_2$-RTC composite map (map9) filtered by local resolution [3].

(*left*) Colored by subunit.

(*right*) Colore by local resolution (key at the bottom).

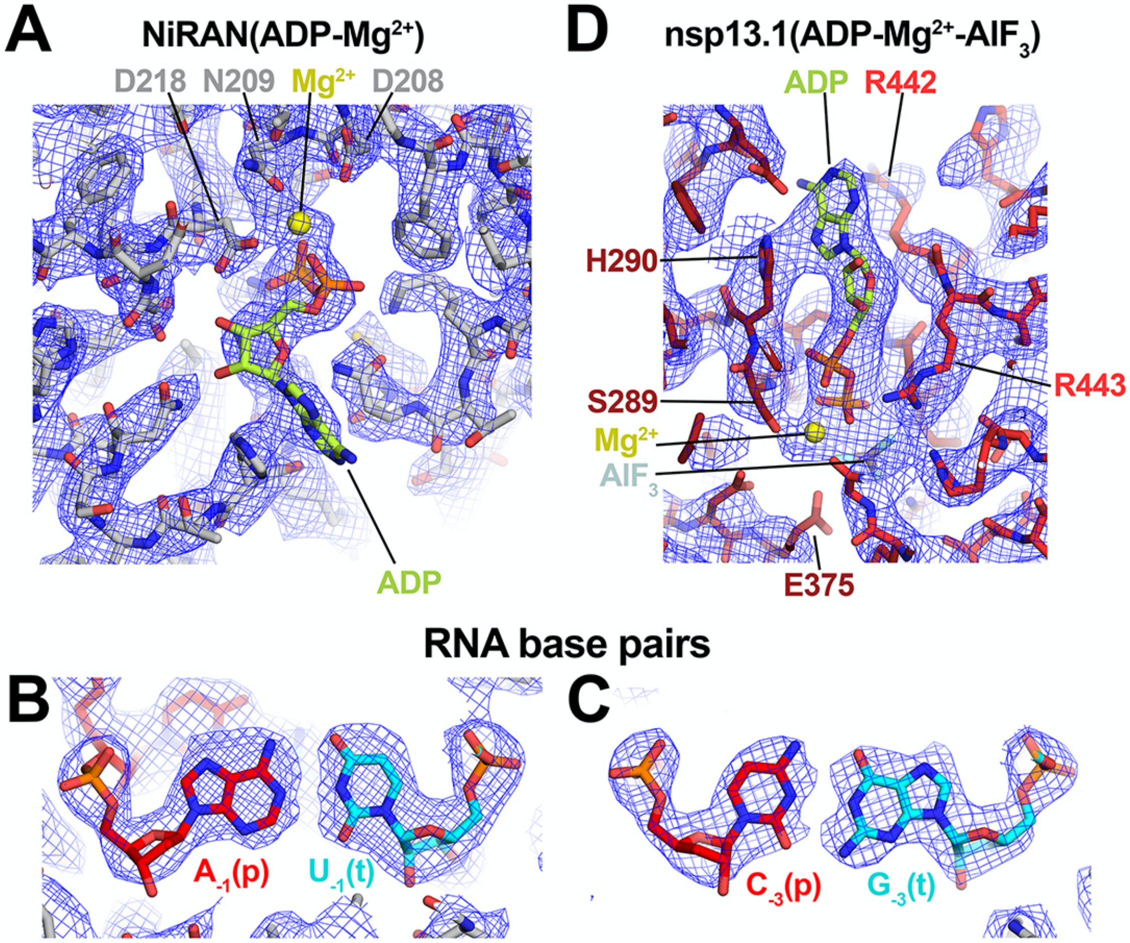

Chen et al., Figure S4

**Fig. S4 | Selected examples of $nsp13_2$-RTC composite cryo-EM map (map9). Related to Fig. 1.**

**A.** NiRAN-ADP-$Mg^{2+}$ bound in the RdRp NiRAN domain.

**B.** AU RNA base pair.

**C.** CG RNA base pair.

**D.** ADP-$Mg^{2+}$-$AlF_3$ bound to nsp13.1.

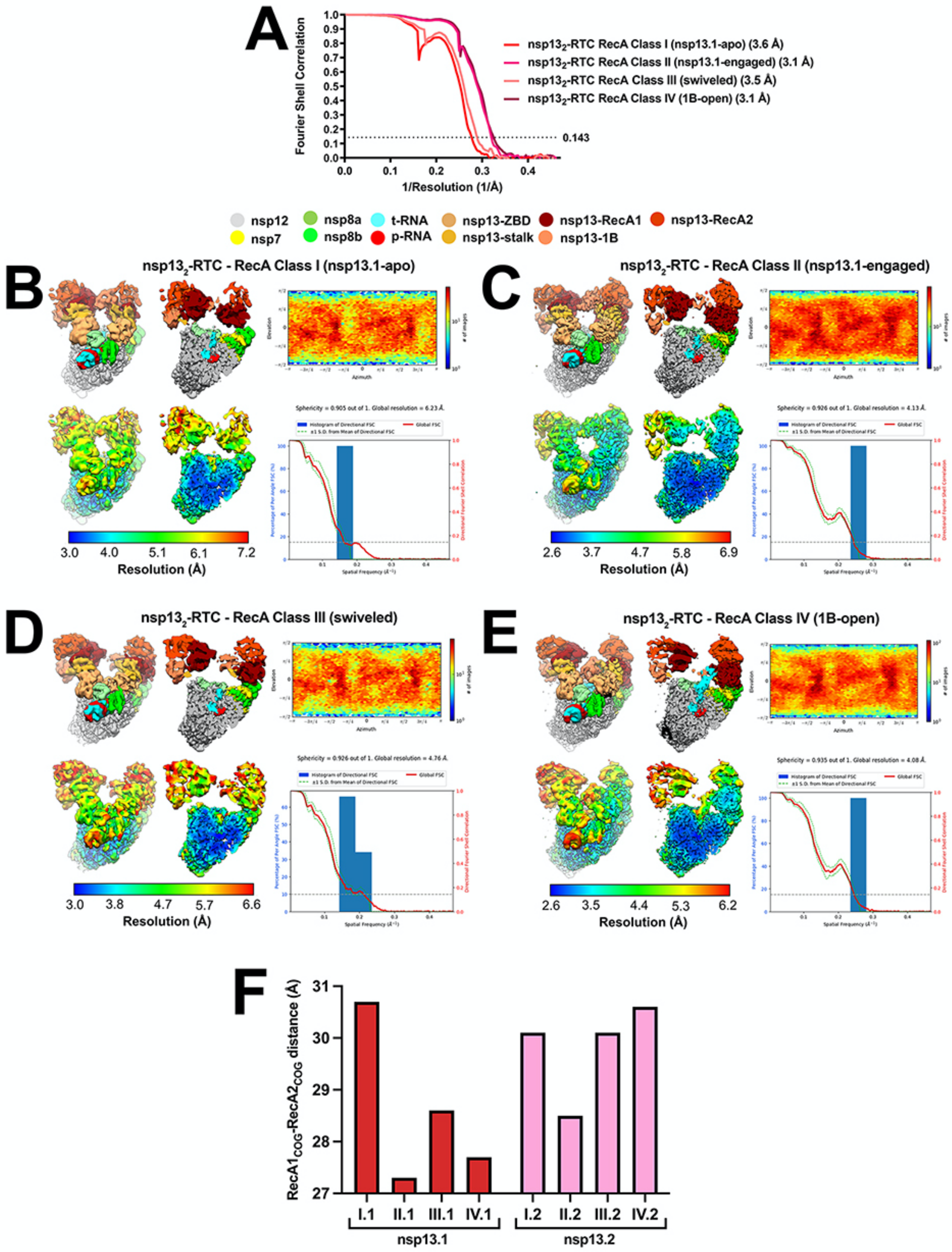

Chen et al., Figure S5

**Fig. S5 | Cryo-EM of $nsp13_2$-RTC classes. Related to Fig. 2.**

**A.** Gold-standard Fourier-shell correlation (FSC) plot for the four $nsp13_2$-RTC classes calculated by comparing the independently determined half-maps from cryoSPARC [5]. The dotted line represents the 0.143 FSC cutoff.

**B - E.** (*left*) The cryo-EM maps filtered by local resolution [3] are shown. The view on the right is a cross-section. In the top row, the maps are colored by subunit according to the key above. In the bottom row, the maps are colored according to the local resolution (color-key indicated in the bar at the bottom).

*(top-right)* Heat map showing particle orientation distribution, calculated using cryoSPARC [5].

*(bottom-right)* Directional 3D Fourier shell correlation calculated by 3DFSC (*top*) [4].

**B.** nsp13.1-apo (map11).

**C.** nsp13.1-engaged (map12).

**D.** Nsp13.1-swiveled (map10).

**E.** 1B-open (map13).

**F.** Histogram denoting the distance separating the RecA1 center-of-gravity ($RecA1_{cog}$) and the $RecA2_{cog}$ for nsp13.1 (red bars, left) and nsp13.2 (pink bars, right) for the four $nsp13_2$-RTC classes.

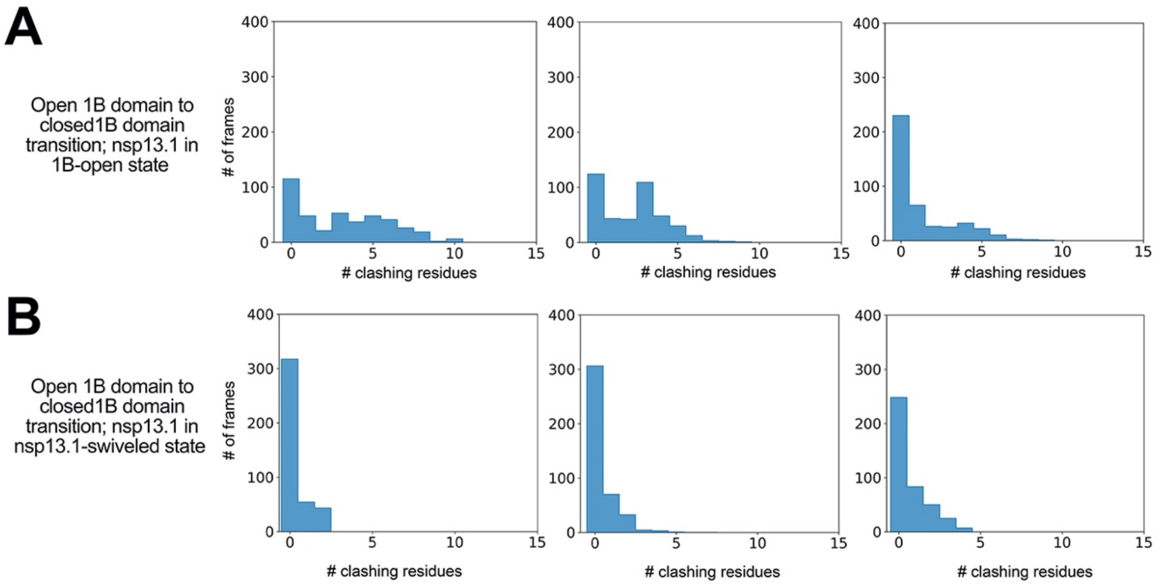

Chen et al., Figure S6

**Fig. S6 | Nsp13.1 1B domain open to closed transition clashes with nsp13.2. Related to Fig. 6.**

The histograms show the number of MD frames in each simulation in which nsp13.1 clashes with nsp13.2 during the nsp13.1-1B domain open to closed states.

**A.** Nsp13.1 in the nsp13.1-engaged position.

**B.** Nsp13.1 in the nsp13.1-swiveled position.

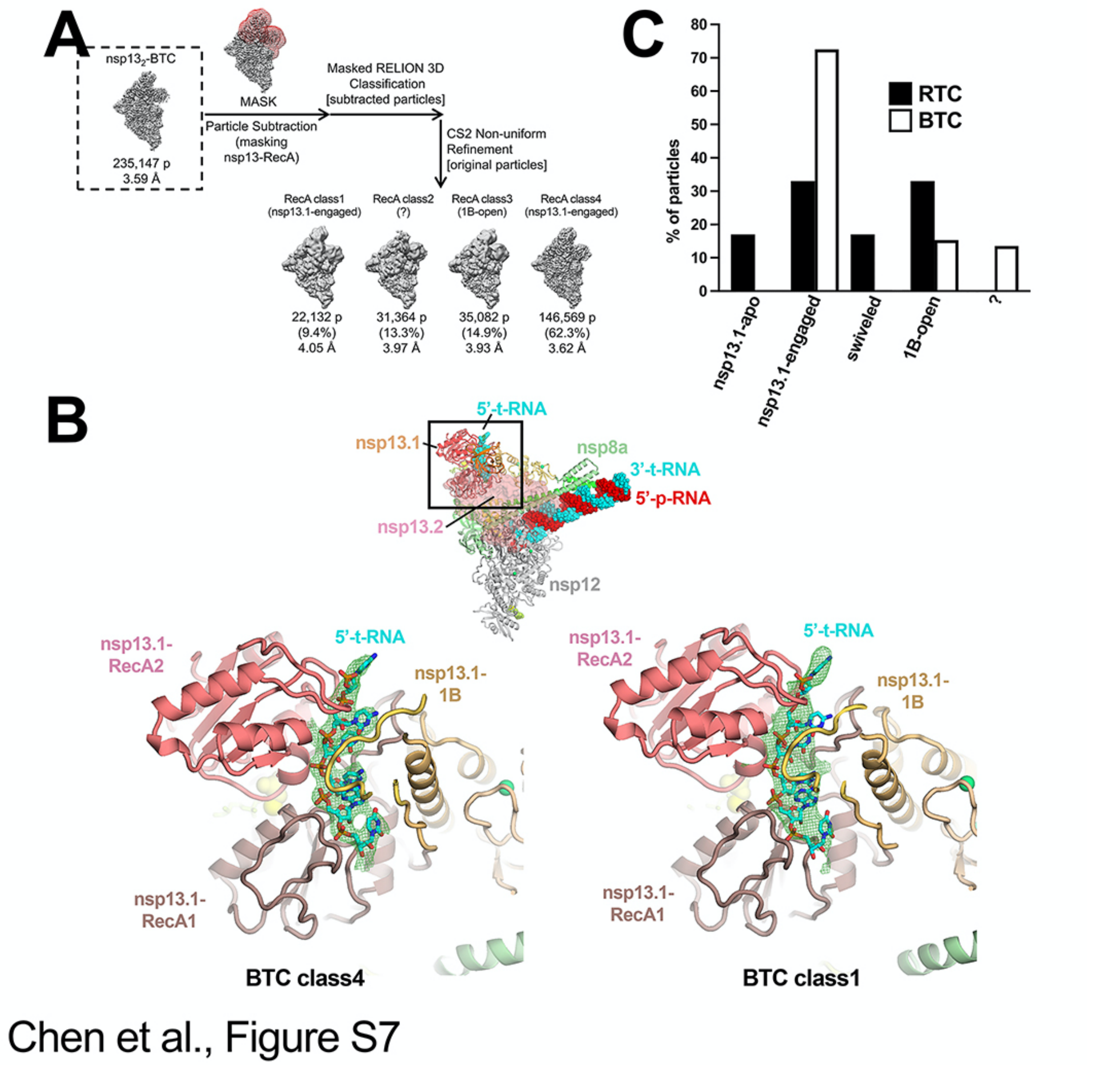

**Fig. S7. Classification pipeline for the nsp13$_2$-BTC dataset and comparison of nsp13$_2$-RTC and nsp13$_2$-BTC class distribution. Related to Fig. 7.**

**A.** Focused classification pipeline for nsp13$_2$-BTC dataset [6].

**B.** (*top-middle*) View of the nsp13$_2$-BTC-class4. Proteins are shown as backbone ribbons except nsp13.2 is shown as a transparent surface. The RNA is shown as spheres. The boxed region is magnified in the views below.

(*bottom*) Magnified views of the nsp13.1 RNA binding channel for nsp13$_2$-BTC-class4 (*left*) and nsp13$_2$-BTC-class1 (*right*). The downstream single-stranded t-RNA is shown as sticks. Cryo-EM difference densities for the downstream single-stranded t-RNA are shown (green mesh).

**C.** Histogram illustrating the nsp13$_2$-RTC particle distribution for the nsp13$_2$-RTC (black bars) and nsp13$_2$-BTC (white bars).

**Supplemental Videos**

**Video S1. Opening of nsp13.1-RecA domains in nsp13.1-apo vs nsp13.1-engaged states. Related to Fig. 4.** The video compares the disposition of the nsp13.1 RecA domains in the nsp13.1-engaged vs. the nsp13.1-apo state. In the nsp13.1-engaged state, the RecA domains are closed onto the substrate RNA and bound to ADP-$AlF_3$, a non-hydrolyzable ATP analog. In the nsp13.1-apo state, the RecA2 domain is rotated open by ~21°, resulting in a translation of the separation between the RecA1 and RecA2 centers-of-gravity by 3.4 Å. The video shows how this conformational change is related to an inchworming model for nsp13.1 translocation, and illustrated how nsp13.1 translocation drives backtracking of the RTC.

**Video S2. Structural overview of the $nsp13_2$-RTC and the nsp13.1-engaged, nsp13.1-open, and nsp13.1-swiveled states. Related to Figs. 2, 3, 5, and 7.** The video highlights the nsp13.1 conformational changes between the nsp13.1-engaged, nsp13.1-swiveled, and nsp13.1-open states.
